# Regulation of kinase activity by combined action of juxtamembrane and C-terminal regions of receptors

**DOI:** 10.1101/2020.10.01.322123

**Authors:** Chi-Chuan Lin, Lukasz Wieteska, Guillaume Poncet-Montange, Kin M. Suen, Stefan T. Arold, Zamal Ahmed, John E. Ladbury

## Abstract

Despite the kinetically-favorable, ATP-rich intracellular environment, the mechanism by which receptor tyrosine kinases (RTKs) repress activation prior to extracellular stimulation is poorly understood. RTKs are activated through a precise sequence of phosphorylation reactions starting with a tyrosine on the activation loop (A-loop) of the intracellular kinase domain (KD). This forms an essential mono-phosphorylated ‘active intermediate’ state on the path to further phosphorylation of the receptor. We show that this state is subjected to stringent control imposed by the peripheral juxtamembrane (JM) and C-terminal tail (CT) regions. This entails interplay between the intermolecular interaction between JM with KD, which stabilizes the asymmetric active KD dimer, and the opposing intramolecular binding of CT to KD. A further control step is provided by the previously unobserved direct binding between JM and CT. Mutations in JM and CT sites that perturb regulation are found in numerous pathologies, revealing novel sites for potential pharmaceutical intervention.

## Introduction

Receptor tyrosine kinases (RTKs) are membrane bound receptors that consist of an extracellular ligand binding domain, a single pass transmembrane region and a cytoplasmic region with kinase activity. Previously it was thought that the initiation of signaling of RTKs required ligand-induced dimerization, followed by a precise order of autophosphorylation on the kinase domain (*1*). However, an increasing number of studies have shown that, in the absence of ligand stimulation, many RTKs are able to self-associate into signaling incompetent dimers (*2*–*6*). Within the context of this unliganded state, the extent of phosphorylation on the kinase domain is restricted to a single tyrosine within the A-loop (*4, 7, 8*) with the exception of EGFR (*9*). This mono-phosphorylated ‘active intermediate’ state is the precursor to the subsequent phosphorylation of additional tyrosine residues. The tight regulation of the active intermediate RTKs in the absence of stimuli is a fundamental precept of numerous cellular outcomes including cell growth, motility, differentiation and metabolism upon ligand stimulation. Dysregulation of the active intermediate state can have devastating effects on ligand-independent signaling, leading to multiple pathologies including cancer, developmental abnormalities and metabolic disorders. Thus, stringent control is required to prevent spontaneous phosphorylation of RTKs within the kinetically favorable, ATP-rich intracellular environment.

Such control is exerted by intracellular amino acid sequences peripheral to KD, both within the juxtamembrane (JM), and the C-terminal tail (CT) regions of the receptor. The modus operandi of these regions varies across different receptors and can lead to both down- and up-regulation of kinase activity (*10*). Structural studies, which have focused largely on the unphosphorylated state, have shown that the binding of JM to KD results in the inhibition of kinase activity of several RTKs (PDGFR (*11*); Eph-family RTKs (*12*); MuSK (*13, 14*); Flt3 (*15*); FGFR1 (*16*); Kit (*16, 17*)). One example is the ephrin receptor B2 (EphB2) in which the JM-KD interaction down-regulates activity through stabilization of the inactive conformation and constraint of the A-loop (*12*). In contrast, the full activity of epidermal growth factor receptor (EGFR) requires the presence of JM which links the asymmetric dimer via a ‘latch’ sequence (*18, 19*). The impact of CT on KD regulation has also been shown to be important in several RTKs. For instance, CT inhibits access of substrates to KD in the unphosphorylated Tie2 receptor (*20*). CT also suppresses the catalytic activity of EGFR through stabilization of an unphosphorylated inactive symmetric dimer (*9, 21*–*24*). The importance of CT in controlling pathogenic signal transduction is demonstrated in the expression of the constitutively active, oncogenic FGFR2 K-*sam*II gene (*25*). There are three variants of K-*sam*II which produce different length truncations of CT (Fig. 1A). Cells in which the truncated K-*sam* gene is amplified exhibit a growth advantage in gastric cancers. Expression of a C3 severely truncated variant in T24 bladder cells leads to un-regulated proliferation (*26*). Thus, the presence of CT prevents proliferative signaling from FGFR2 through an imprecisely known mechanism.

**Fig. 1.**
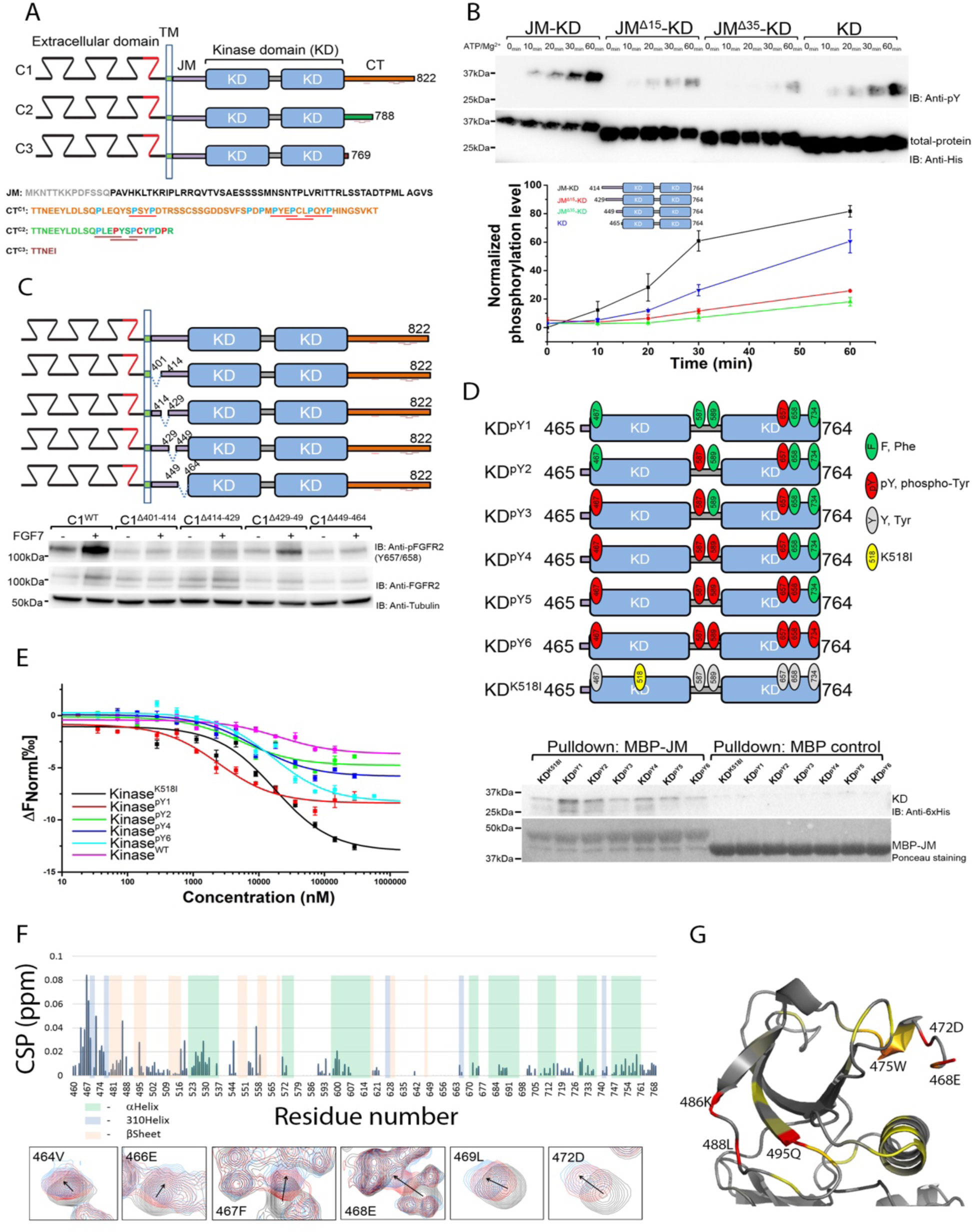
The presence of JM enhances activity of KD. (**A**) Schematic diagram of FGFR2IIIb C1, C2, and C3 isoform. These isoforms differ only in their C-terminal tail. C1 isoform (orange) includes the full length C-terminal tail (CT^C1^); C2 isoform (green) has a truncated C-terminal tail (CT^C2^) and also includes three point mutations (red); the C3 isoform (purple) is truncated by 56 residues (CT^C3^). Proline residues are shown (blue) and proline-rich motifs in C1 and C2 isoforms are underlined. JM sequence is also shown (black) – membrane-bound sequence of JM (grey). (**B**) *In vitro* kinase assay using progressive JM-deletions in JM-KD (residues 414-764, black; 431-764, red; 447-764, green; and 465-764, blue). 100 nM of each protein was used for the *in vitro* phosphorylation assay (see STAR Methods for details). Phosphorylation levels were determined using a general pY99 antibody. His-tag antibody was used for total protein control. (**C**) Intermittent deletions in JM down-regulate FGFR2 activity. HEK293T cells were transfected with FGFR2 with JM deletions (C1^Δ401-414^, C1^Δ414-429^, C1^Δ429-449^, and C1^Δ449-464^). Cells were serum starved overnight and left unstimulated or stimulated with 10ng/ml FGF7 for 15 min. Cell lysates were blotted with indicated antibodies. (**D**) Binding of JM to progressively phosphorylated KD. Six tyrosine residues on KD were mutated to mimic the sequential phosphorylation pattern of KD (KD^pY1^ to KD^pY6^; fig. S1A). MBP-JM was used to pulldown KDs (His-tagged). (**E**) MST measurements of the binding affinity between JM and KD with different phosphorylation levels. JM was labelled with Atto 488 dye and serial dilutions of KD were titrated at 25°C. (**F**) CSPs of ^15^N-KD^pY1^ titrated by JM derived from ^1^H-^15^N HSQC spectra (fig. S1E). Large changes occur on the N-terminal lobe. Selected residues experiencing major CSPs are shown as the bottom panels. (**G**) CSPs from residues in ^15^N-KD^pY1^ as JM is titrated plotted on the structure of KD^pY1^. CSPs shown as gradient: Yellow – lower, Red - higher. Selected residues with major CSPs are labelled.

Our current knowledge of the regulatory roles of JM and CT is restricted to experiments based on the unphosphorylated KD with either JM or CT independently. The potential for cooperation between JM and CT towards regulation of the mono-phosphorylated, active intermediate state of the KD in the absence of RTK stimulation remains unexplored. FGFR2 provides a good example of a highly regulated RTK, particularly since the A-loop Y657-mono-phosphorylated state prevails under unliganded conditions (*4*). In this state FGFR2 is primed to respond rapidly to growth factor binding to produce the phosphorylated platform for recruitment of downstream effector proteins, but is subjected to stringent controls. Here we reveal the intricate mechanism by which the interplay of JM and proline-rich sequences on CT enable the receptor to sustain the active intermediate under non-stimulated conditions, and yet inhibit further catalytic activity. Our data also provide a rationale for the uncontrolled proliferation of the K-*sam* C-terminally truncated variants. Since virtually all RTKs possess JM and proline-rich sequences within their CT regions, this fine-tuning of regulation by JM and CT is likely to be conserved across RTKs. Indeed, mutations of proline residues on numerous RTK CTs are connected to dysregulation of kinase activity (*27*–*29*) leading to a range of human pathologies.

## Results

### Activity of KD is enhanced in the presence of JM

JM of FGFR family receptors was predicted to play an autoinhibitory role on KD activation (*16*), however, direct evidence of this is lacking. The impact of JM (residues 414-465) on kinase function was investigated through four dephosphorylated FGFR2 JM-KD constructs with progressively increasing truncations of JM. The rate of phosphorylation of KD was greater with intact JM (Fig. 1B). However, deletion of the entire JM resulted in phosphorylated product, as would be expected for an unencumbered enzyme in solution. The influence of JM on phosphorylation was measured in HEK293T cells over-expressing full length FGFR2IIIb (KGFR/K-*sam*-IIC1, C1 isoform, FGFR2^C1^) including short, intermittent JM fragment deletions (Fig. 1C). Immunoblotting the A-loop phosphorylated tyrosines (pY657/pY658) in both basal and FGF7-stimulated phosphorylation of FGFR2^C1^ was significantly reduced in all JM deletion variants confirming the importance of the intact JM.

To understand the precise regulatory function associated with JM to KD interaction in FGFR2, we sought to understand the effect of progressive phosphorylation of KD on JM binding. Direct interaction between an MBP-JM and a series of six mutants that mimic the sequential phosphorylation pattern of KD (KD^pY1^ to KD^pY6^; schematic Fig. 1D and fig. S1A) revealed that JM bound most strongly to the mono-phosphorylated KD^pY1^ (i.e. the ‘active intermediate’ state. Fig. 1d. K_d,app_ = 2.51 ± 0.20 µM; Table S1 and Fig. 1E and fig. S1B). The affinity of JM for KD reduced with progressive phosphorylation. Only weak binding was apparent with the unphosphorylated, catalytically inactive K518I mutant, KD^K518I^.

To determine the precise region through which JM and KD interact, we generated five JM peptides of 15-16 amino acids and measured their affinities to KD^pY1^ (table S1 and fig. S1C). ^414^PAVHKLTKRIPLRRQVT^430^ demonstrated the tightest binding (K_d_ = 36.8 ± 6.2 µM) while ^407^KPDFSSQPAVHKLT^420^ bound only three-fold weaker. These peptides share the consensus sequence ^414^PAVHKLT^420^ proximal to the N-terminal of JM.

KD residues that interact with JM were identified by NMR. Titration of JM into ^15^N-labelled KD^pY1^ led to major chemical shift perturbations (CSPs) in the N-lobe of KD^pY1^ resulting from changes in chemical environment on binding; including residues 464-472 (Fig. 1F) as well as 486 and 495 (Fig. 1G) (see fig. S1D for the assignment coverage and fig. S1E for HSQC titration spectrum). Notable CSPs were also observed in residues located within the αC helix (525-530; fig. S1F), which is a dynamic regulatory element of the KD. This suggests a mechanism whereby the binding of JM to the intermediate KD^pY1^ state would promote further kinase activity.

### JM binding enhances asymmetric KD dimer formation

Asymmetric dimerization is crucial for receptor enzymatic function. Having shown that binding of JM is important in promoting kinase activity, we next investigated the impact of JM on dimerization. A series of JM-KD polypeptides exhibiting progressively increasing phosphorylation states (fig. S2A and S2B) exhibited the highest dimer population in the mono-phosphorylated state, JM-KD^pY1^ (Fig. 2A). In both the catalytically inactive mutants JM-KD^K518I^ and JM-KD^Y657/658F^, dimerization was abrogated. The dependence of both JM binding and dimerization on the phosphorylation state of KD is consistent with JM acting as an intermolecular latch which is released with increasing pY burden.

**Fig. 2.**
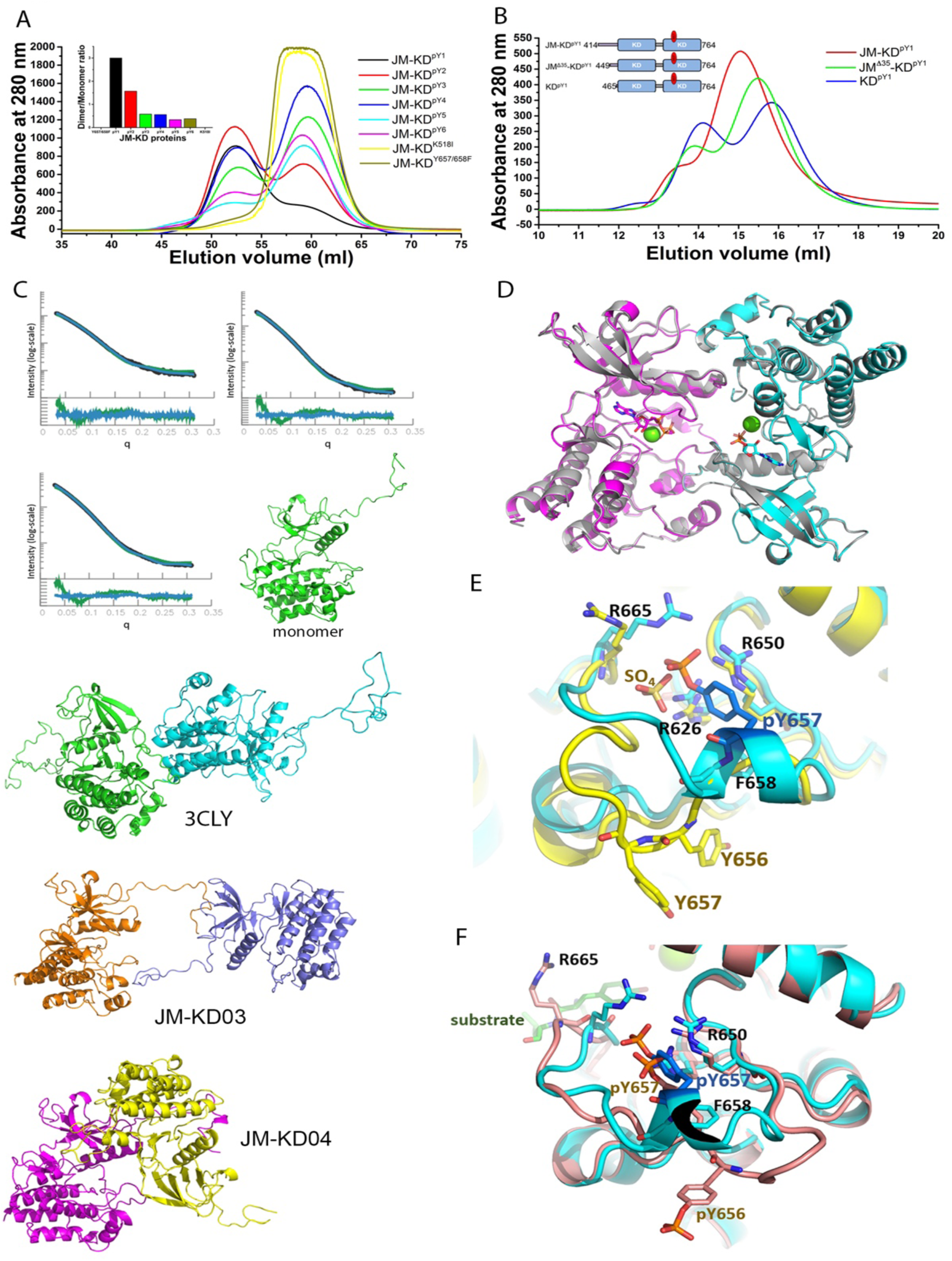
JM forms an intermolecular interaction in an asymmetric KD dimer. (**A**) JM-KD construct with progressively increasing pY residues on KD were run on size exclusion chromatography (SEC) at 60 - 80 µM (inset: dimer/monomer ratio of KD with different phosphorylation levels). (**B**) Dimerization of KD^pY1^ is reduced in the presence of JM. The dimerization of JM-KD^pY1^ constructs with JM deletions (JM-KD^pY1^; JM^Δ35^-KD^pY1^; and KD^pY1^) was determined using 10 µM injected on a size exclusion column. (**C**) SAXS description of the structure of JM-KD^pY1^. Top panel: the calculated SAXS pattern for 1-state (green) and 3-state (blue) models fitted to the experimental data (black) at different total KD concentrations (left plot, 65 µM; middle, 130 µM and right 210 µM); bottom panel: residuals of fit for 1-state (green) and 3-state models (blue). The resolution, q, is given in 1/Å and the intensity is given in arbitrary units. Model structures based on crystal structure-derived and *in silico*-docked dimeric and multimeric assemblies. Mono – structure of the monomeric state, JM-KD03 - two kinases (orange/blue) loosely connected through their JM interactions; JM-KD04 asymmetric dimer where JM of the enzyme-like state (magenta) binds to the substrate-and cyan). **(D**) Asymmetric unit of KD^pY1^ containing four molecules. Phosphorylated chains A (magenta) and C (cyan) superimposed onto chains C and D (grey). ATP shown as stick model, and Mg^2+^ as green sphere. (**E**) A-loop superposition between KD^pY1^ (chain B, cyan) and unphosphorylated kinase with 2PSQ (chain A, yellow). **(F**) Zoom into the A-loop between KD^pY1^ (chain B) and dephosphorylated kinase with 2PVF (salmon).

To establish further how the JM affects the dimeric state of KD^pY1^ we used three JM-KD^pY1^ constructs with progressively truncated JM (schematic Fig. 2B). Truncation of JM resulted in an increase in the population of dimers, with isolated KD^pY1^ showing the greatest population of dimer. We quantified the ‘apparent’ dimerization constant of the mono-phosphorylated KD in the absence of JM (KD^pY1^: K_d,app_ = 112 ± 9 nM; fig. S2C). When JM was present the dimerization affinity was reduced by an order of magnitude (JM-KD^pY1^: K_d,app_ = 3.46 ± 0.10 µM by MST (fig. S2C) and K_d,app_ = 3.07 µM by surface plasmon resonance, SPR (fig. S2D)). This, somewhat counter-intuitive result, suggests that although the presence of JM increases phosphorylation, it weakens dimerization of the KD. It also confirms that KD^pY1^ and JM-KD^pY1^ dimers are conformationally different.

This was confirmed using small angle X-ray scattering (SAXS). In the absence of JM, KD^pY1^ scattering consistently revealed equilibrium between monomers, symmetric head-to-tail dimers and additional larger particles, possibly residual non-specific higher-order forms (fig. S2E and table S2). The symmetric dimer corresponds to that depicted in crystal structures of inactive FGFR2 (PBD 2PSQ (*30*); KD05 in fig. S2E) where access to the active site, and the positioning of the A-loop of one molecule are all encumbered by the presence of the second molecule. In stark contrast to KD^pY1^, in the selected ensembles of the extended JM-KD^pY1^ the symmetric head-to-tail dimer was replaced by asymmetric dimers, with ∼ 10% of these loosely connected through JM (Fig. 2C). This dramatic redistribution of polypeptide conformations is consistent with that required to produce the reduction in dimer affinity observed for JM-KD^pY1^ compared to KD^pY1^ (fig. S2C). The structures of these dimers are reminiscent of reported structures of the asymmetric conformations of enzyme-like (aka. receiver) and substrate-like (aka. activator) KDs caught in the act of trans-phosphorylation (*30, 31*). In these modelled dimeric structures, JM from one protomer binds to the other protomer, leaving one JM unoccupied (fig. S2F). Collectively, our data suggest that in the absence of ligand stimulation, through forming a latch which stabilizes the asymmetric conformation between mono-phosphorylated KDs, and hence abrogating symmetric dimer formation, JM is able to sustain a dynamic relationship between mono-phosphorylated KDs from two FGFR2 molecules to promote kinase activity and facilitate access to substrate sites. This then requires that additional control mechanism(s) have to be in place to prevent unrestrained phosphorylation of the active intermediate.

Further detail of KD^pY1^ was provided by its 2.5 Å crystal structure determined in complex with a non-hydrolysable ATP analogue (PDB ID 6V6Q, table S3). The asymmetric unit contained four molecules which were arranged in two symmetric head-to-tail dimers (Fig. 2D and fig. S2G). These dimers correspond to the symmetric dimers observed in SAXS (fig. S2E). Two of the four molecules in the asymmetric unit showed well-defined electron density for the A-loop clearly featuring pY657 (fig. S2H). pY657 is positioned within a single turn α-helix, and the phosphate is coordinated by three arginine residues: R626, R650 and R665 (Fig. 2E). Burial of pY657 in this way has important consequences for A-loop mobility and sustaining the mono-phosphorylated state. In support of our SAXS data, the dimeric juxtaposition of KD^pY1^ is the same as that adopted in the published unphosphorylated kinase structure (PDB 2PSQ; Fig. 2D and fig. S2G) even though 2PSQ crystals do not contain ATP and form a different crystal lattice. The only notable differences between the two structures are found in the A- and nucleotide-binding loops. In contrast to our structure, the A-loop tyrosine residues Y656 and Y657 (corresponding to Y657 and Y658 in FGFR2IIIb) are solvent exposed in 2PSQ (Fig. 2E). KD^pY1^ also shows features similar to the dually phosphorylated active substrate-bound kinase structure (PDB 2PVF; fig. S2I). The orientation of N and C-lobes, the αC helices, and the catalytic residues K518 and E535 in the KD^pY1^ structure are superimposable with 2PVF. However, the A-loop in 2PVF does not fold into a helix around pY657, but instead is linear (Fig. 2F). We concluded that in the absence of JM, KD^pY1^ forms a symmetric dimer which imposes constraint on the A-loop resulting in occlusion of access of substrate and corrupts the site for correct placement of the γ-phosphate of ATP in the active site. The symmetric KD dimer structure is potentially relevant to the unphosphorylated receptor or when KD is constrained as part of the GRB2-bound heterotetramer (*4*). Furthermore, the engulfment of pY657 in the kinase core provides a possible mechanism to preserve the active intermediate mono-phosphorylated state.

To determine whether JM from the enzyme-like, or substrate-like protomer forms the latch and facilitates trans-phosphorylation, we incubated catalytically inactive KD^K518I^ (the substrate) with JM-KD^pY1^ or KD^pY1^ (the enzymes). In this case the presence of JM increased phosphorylation of KD^K518^ (fig. S2J). We then incubated KD^pY1^ (the enzyme) with JM-KD^K518I^ or KD^K518I^ (the substrates) and observed no difference phosphorylation of the two substrates (fig. S2K). Deletion of ^420^TKR^423^ and ^426^RRQ^428^ on the enzyme-like JM-KD^pY1^ (within the ^414^PAVHKLTKRIPLRRQVT^430^ KD^pY1^-binding sequence), had the biggest impact on down-regulating phosphorylation of KD^K518I^ (fig. S2L). Our data show that in the active intermediate state JM from the enzyme-like protomer latches on to the substrate-like protomer in the asymmetric active dimer and increases the phosphorylation of the latter molecule. In the absence of other regulatory interactions, this latch holds the active KDs such that they can interact with one another, whilst being prevented from self-association into the inactive symmetric dimer.

### Activity of KD is inhibited by CT

To investigate the detailed regulatory function of CT, we first measured the impact on receptor activity of N-terminally Flag-tagged, FGFR2IIIb C-terminally truncated C1, C2, C3 K-*sam*II isoforms. HEK293T cells transfected with FGFR2^C1^; FGFR2^C2^; FGFR2^C3^ or FGFR2^C1Δ34^ (FGFR2^C1^ with 34 amino acids deleted from the C-terminus that is identical in length to C2 but does not contain the Q779P, S783C and T787P mutations which produce a sequence of two consecutive PXXP motifs: CT sequences shown in Fig. 3A) revealed that deletion of CT led to increased receptor phosphorylation and activation of effector proteins under both unliganded and FGF7-stimulated conditions (Fig. 3A, FGFR2^C1Δ34^). The absence of CT in FGFR2^C3^ promoted downstream signalling in both ERK1/2 (MAPK) and AKT pathways without ligand stimulation. This is likely to be due to the binding of the scaffold protein FRS2 which is known to bind to the JM and mediate downstream effector protein recruitment to the activated receptor (Fig. 3A) (*32*) (in contrast to FGFR1 to which FRS2 is constitutively bound; (*33*)). This result strongly suggests that the presence of CT controls FGFR2 kinase activity as well as inhibiting the interaction of FRS2 with the receptor.

**Fig. 3.**
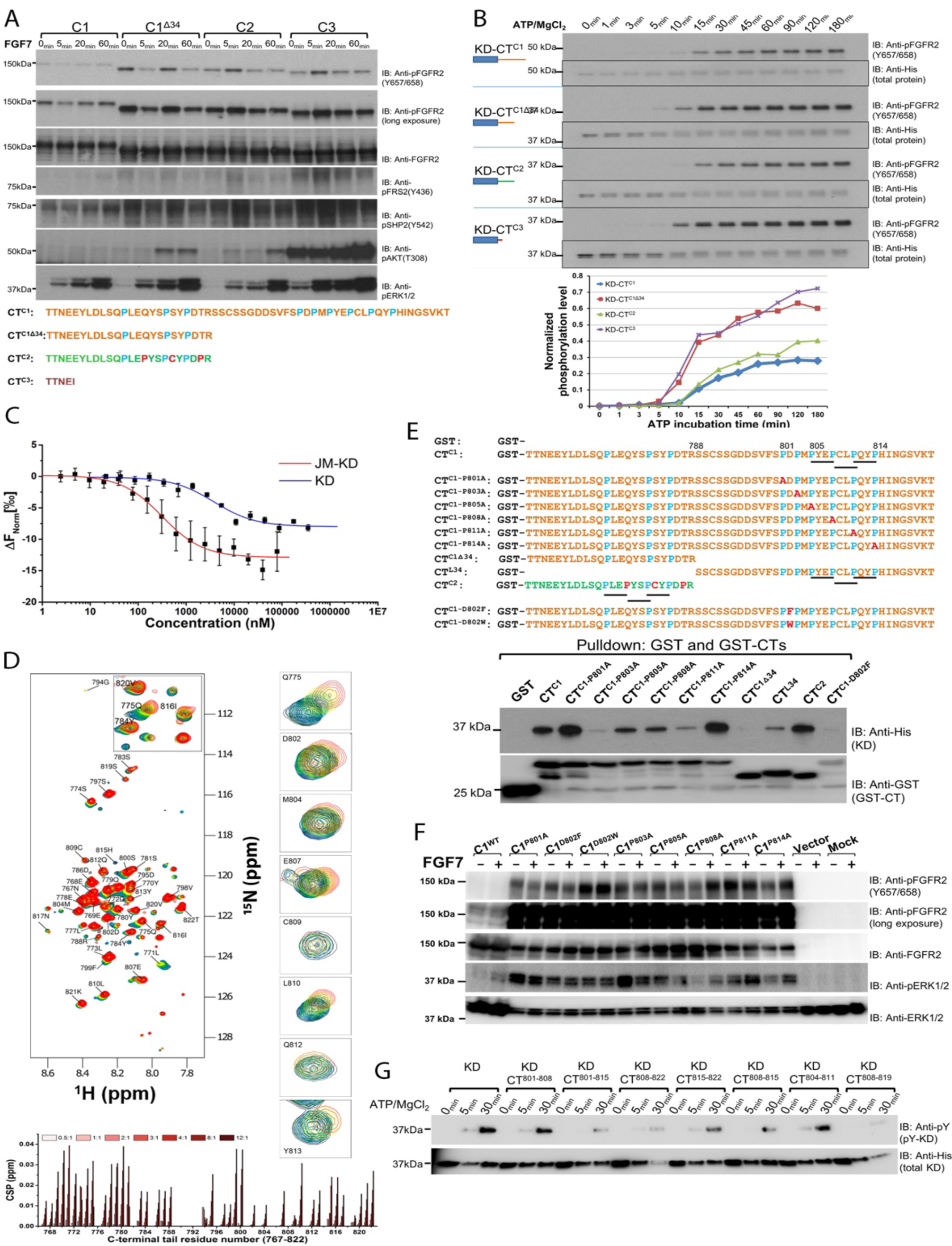
Proline-rich motifs interact and downregulate kinase activity. (**A**) Immunoblotting analysis of signalling activity of FGFR2IIIb isoforms. FGFR2^C1^; FGFR2^C1Δ34^; FGFR2^C2^ and FGFR2^C3^ were transfected into HEK293T cells. Cells were starved or stimulated with 10ng/ml FGF7. The levels of receptor phosphorylation and downstream activity on each isoform were probed with the indicated antibodies. (**B**) Proline-rich CT inhibits *in vitro* kinase activity. Recombinant KD-CT^C1^, KD-CT^C2^, KD-CT^C3^ and control clone: KD-CT^C1Δ34^ were incubated with ATP/Mg^2+^ at 25 °C and quenched with 100 mM EDTA at different time points as indicated. The activation level was measured using an anti-pY657/658 antibody. Bottom: The densitometry analysis of kinase activity (KD-CT^C1^ – blue; KD-CT^C2^ – green; KD-CT^C3^ – purple and KD-CT^C1Δ34^ – red). (**C**) The affinities of CT^C1^ to JM-KD^pY1^ (red) and KD^pY1^ (blue) determined using MST. CT^C1^ was labelled with Atto488 dye then titrated with unlabelled JM-KD^pY1^ and KD^pY1^. (**D**) HSQC spectra of unbound ^1^H-^15^N-labelled CT^C1^ overlaid with KD^pY1^-bound CT^C1^ at different ratio (black (0:1) to red (12:1)). Examples of peaks with high chemical-shift perturbations (CSPs) are shown by labels indicating the assignment of given peaks. CSPs chat of ^15^N-KD^pY1^ titrated by CT^C1^ was derived from ^1^H-^15^N HSQC spectra. Large changes occur on both N-terminal and C-terminal residues of CT^C1^. (**E**) Wild type GST-CT^C1^ and its individual P to A mutants, the first 24 residues of CT (CT^C1Δ34^), the last 34 residues (CT^L34^), CT^C2^ and CT^C1^ D802F or D802W (to explore the importance of the charged acid group in binding) were used for a GST pulldown experiment with KD^pY1^. The symmetric dimerization of KD^pY1^ at the concentration range used in this experiment (1 µM) was assumed to have negligible impact on binding of the various CT variants. (**F**) The presence of the intact proline-rich motif in FGFR2^C1^ inhibits both FGFR2 and downstream ERK1/2 activities. FGFR2^C1^ variants with individual P/A, D/F, and D/W mutants as indicated were transfected into HEK293T cells. Cells were starved or stimulated with 10ng/ml FGF7 for 15 minutes. Cell lysates were blotted with indicated antibodies to examine the importance of the proline-rich sequence on CT. (**G**) Dephosphorylated KD was incubated with seven CT-derived short peptides including fragments of the proline-rich sequences (residues 801-808; 801-815; 808-822; 815-822; 808-815; 804-811 and 808-819) to test their ability to regulate kinase activity. See STAR Methods for phosphorylation and quenching procedures.

We next investigated the direct effect of CT on the regulation of FGFR2 kinase activity using recombinant protein. Consistent with the cell-based assay (Fig. 3A), the phosphorylation of the A-loop increased as CT was truncated (KD-CT^C3^ Fig. 3B). Thus, KD-CT^C3^, like KD^pY1^ which appeared in dynamic equilibrium between monomers and symmetric head-to-tail dimers (Fig. 2B, 2D and fig. S2E), behaved as a free enzyme (Fig. 1B). As expected, KD-CT^C1^ had the lowest kinase activity but CT in KD-CT^C1Δ34^ released inhibition as seen in our cell-based assay. However, KD-CT^C2^, which is of an identical length, but includes similar PXXP motifs as present in KD-CT^C1^, restored the inhibitory capability. This result suggests the importance of the PXXP motif(s) of CT in the inhibition of kinase activity.

### A proline-rich motif on CT is required for the binding to KD

So far, our data indicate that when CT is present it inhibits the active intermediate receptor from progressing to the fully phosphorylated state. This was hypothesized to occur via two distinct mechanisms; 1) antagonistically blocking receptor activation through direct binding to KD, and/or 2) through binding of CT to KD and/or JM to inhibit formation of asymmetric dimer.

CT^C1^ bound to KD^pY1^ with a moderate affinity (K_d_ = 3.75 ± 0.46 µM; Fig. 3C). Although KD^pY1^ is potentially in the form of a dimer, the profile of the binding curves suggests that CT binds independently to the domain. NMR spectroscopy was used to probe the interaction surfaces of CT^C1^. To this end, CSPs of ^15^N-labelled CT^C1^ were measured on addition of KD^pY1^ (Fig. 3D). Two distinct potential interacting regions of CT^C1^ were observed; residues around 765 to 780, including the known binding site for downstream effector proteins Y770 (Fig. 3D), and residues within the proline-rich motif in the C-terminus (D802 to Y813: Fig. 3D). In a separate experiment CT^C1^ was divided into two fragments: the first 24 residues 765-788 (CT^C1Δ34^) and the last 23 residues 800-822 (CT^L23^). CT^C1Δ34^ revealed negligible CSPs changes (fig. S3A and S3B) which supported our previous kinase assay data which showed that in the absence of the PXXP motifs this region does not affect kinase activity. More significant CSP changes from CT^L23^ were observed (fig. S3C and S3D). We concluded that the last 23 residues containing the PXXP motifs are necessary for binding with the KD^pY1^, and this facilitates subsequent engagement of the first 24 residues of CT^C1^. CSP and affinity measurements of peptide fragments of CT revealed that the tightest binding sequence was ^801^PDPMPYEPCLPQYPH^815^ (K_d_ = 25.9 ± 5.4 µM; table S1 and fig. S3E). Our NMR experiments confirm that although a potentially extensive interface is involved, the PXXP motifs are required for CT^C1^ to interact with the active intermediate KD^pY1^ and hence inhibit kinase activity.

The importance of individual proline residues within the PXXP motif in binding to KD^pY1^ was investigated using an *in vitro* GST pulldown assay with both the GST-tagged CT^C1^ and CT^C2^ (Fig. 3E). Binding was significantly reduced in the presence of proline to alanine point mutations except for P801A and P814A. The mutation of both P803 and D802 had a large impact on binding. Supported by our kinase assay and NMR data, we found that GST-CT^C2^ bound to KD^pY1^ whilst the first 24 amino acids (GST-CT^C1Δ34^) of C1 did not. The similarity of CT^C2^ with the wild type CT^C1^ was also apparent in the kinase phosphorylation data (Fig. 3B). Sequence alignment suggests that the interactions are strongest when the sequence includes a PXEPXXPXYP motif (where X is any residue) which occurs between residues 805 and 814 for CT^C1^ and 776 and 785 for CT^C2^.

Point mutations in FGFR2^C1^ PDPMPXEPXXPXYP sequence confirmed the importance of this region for signaling in HEK293T cells (Fig. 3F). Even in the absence of FGF7 the corruption of the proline-rich sequence had a dramatic affect in up-regulation of FGFR2 and its downstream ERK1/2 signalling. Inhibition of recombinant KD by incubating peptides derived from CT^L23^ identified ^808^PCLPQYPH^815^ as the minimum sequence of CT required for KD down-regulation (Fig. 3G and table S1 for sequences).

### The CT-KD interface includes regions associated with kinase activity

Having established a detailed knowledge of the residues from CT that interact with KD^pY1^, we also measured CSP data for ^15^N-labelled KD^pY1^ on addition of CT. (Fig. 4A). Mapping the perturbed residues onto the crystal structure (Fig. 4B) revealed four clusters of residues in the KD^pY1^ that showed significant shifts: 470-490 at the very N-terminal end of the N-lobe (including the nucleotide binding loop); 515-525 (including the regulatory αC helix); 590-610 (including the kinase insert), and 710-740 at the very C-terminal end of the C-lobe. The interface between CT^C1^ and KD^pY1^ is therefore extensive and covers regions associated with kinase activity providing a rationale for the inhibitory nature of the direct interaction. Comparing CSPs for binding of JM to KD^pY1^ (Fig. 1F) and CT to KD^pY1^ (Fig. 4A) suggests that there is some overlap, although residues with the largest CSPs are not entirely coincident. We performed HSQC titration of JM with ^15^N-labelled KD^pY1^ and recorded the CSPs for residues in regions previously seen to interact with JM on KD^pY1^ (464V, 466E, and 467F at the N-terminal lobe and 530S and 532L on the αC helix: fig. S4A, left panels). Subsequent titration of CT to the preformed JM-KD complex resulted in negligible CSP effects on the JM-bound KD^pY1^ spectra (fig. S4A, right panels) underscoring that binding of CT does not compete directly with JM for binding to KD^pY1^.

**Fig. 4.**
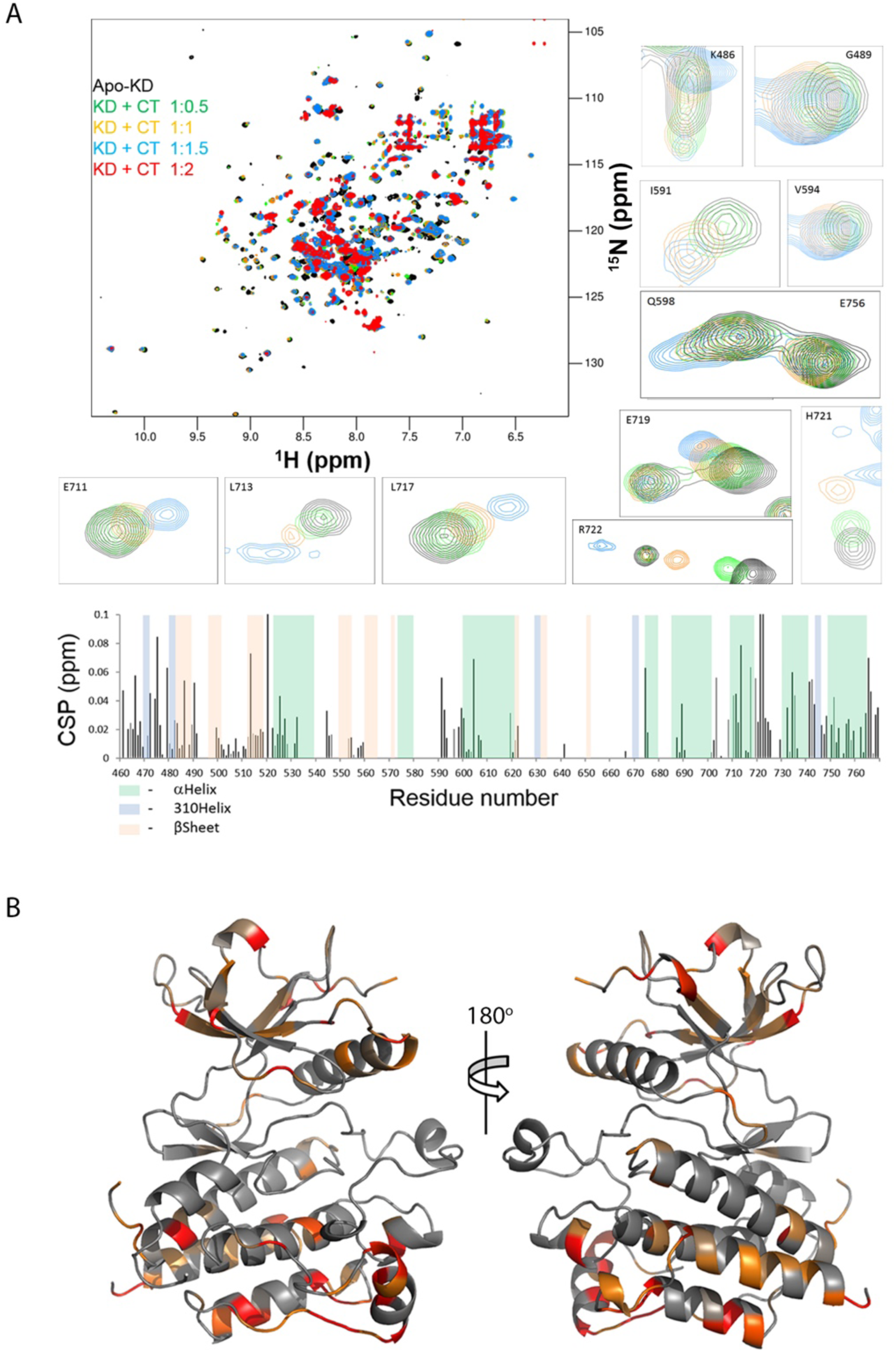
Identification of CT binding site on KD using NMR. (**A**) ^1^H-^15^N HSQC spectra of unbound FGFR2 KD^pY1^ (Black) overlaid with CT^C1^ bound KD^pY1^ at different concentration ratios. Examples of peaks with major CSPs shown. CSPs chart of CT^C1^ titrated into ^15^N-KD^pY1^ derived from ^1^H-^15^N HSQC spectra. Large CSPs are observed at both N-terminal and C-terminal lobes. (**B**) CSPs of CT^C1^ binding to KD^pY1^ plotted on the X-ray structure of mono-phosphorylated kinase. As shown in Fig. 1G, a red-to-yellow gradient was used to indicate the CSP residues. Minor CSPs appear on one side of the regulatory αC helix, suggesting the binding of JM could regulate kinase activity.

### CT inhibits asymmetric dimerization of JM-KD^**pY1**^

When JM is present with KD^pY1^ an asymmetric dimer forms which can promote further kinase activity. Conversely the presence of CT with KD inhibits this activity. We therefore sought to understand how these opposing control mechanisms combine to regulate further activity of KD^pY1^. Using size exclusion chromatography, we showed that CT was able to block dimer formation when included as part of KD^pY1^-CT (Fig. 5A). GST-CT^C1^ was able to pull down (Fig. 5B) and form a high affinity complex with JM-KD^pY1^ (SPR, K_d_ = 165.2 ± 1.6 nM; MST, K_d_ = 304 ± 44 nM: table S1, Fig. 3C and fig. S5B respectively). The binding of CT to JM-KD^pY1^ was able to disrupt the asymmetric dimerization of the construct, as demonstrated using steady-state fluorescence resonance energy transfer (FRET) measurement (fig. S5A). Increasing the phosphorylation state of KD reduced the affinity of CT (fig. S5B). CT can bind to the non-phosphorylated JM-KD^Y657/658F^ but with reduced affinity (K_d_ = 5.65 ± 1.9 µM; table S1). It is notable that since the binding of CT to JM-KD^pY1^ disrupts the asymmetric dimerization of the latter, the resulting high affinity complex has molecular ratio CT:JM-KD^pY1^ of 1:1.

**Fig. 5.**
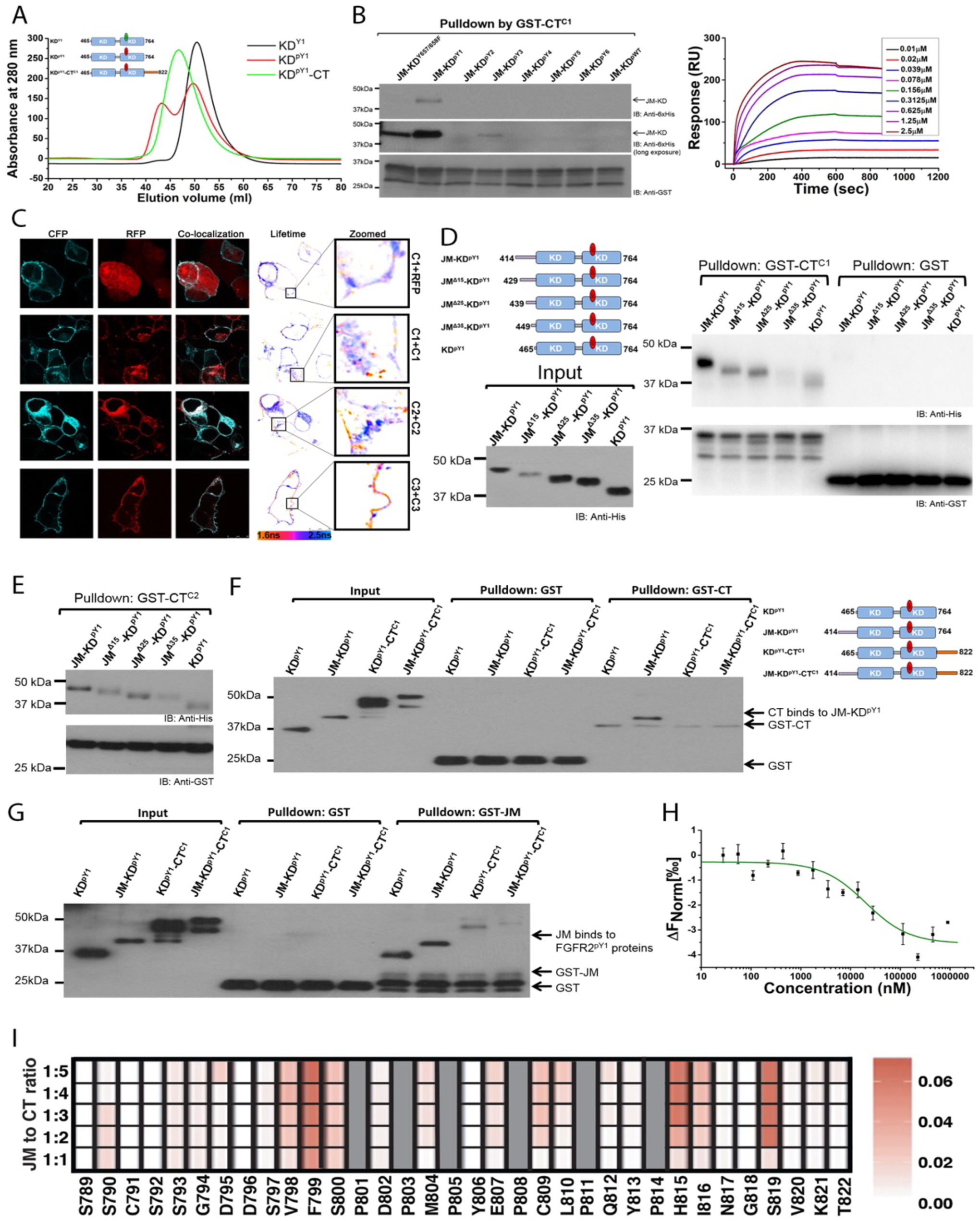
CT binding to KD^pY1^ disrupts the formation of asymmetric dimer. (**A**) Dimerization status of FGFR2 kinases in the presence or absence of CT. KD^pY1^ (blue curve) includes a dominant population of dimers at 60 µM in SEC, whereas the unphosphorylated KD (green curve) prevails as a monomer. The mono-phosphorylated KD^pY1^-CT construct (red curve) also exists as a monomer in solution. (**B**) Phosphorylation of JM-KD affects CT binding. GST-CT^C1^ was used to pulldown the progressively increasing phosphorylation of JM-KD constructs. The pulldown indicates that JM-KD^pY1^ was the best binding partner for GST-CT^C1^. (**C**) Dimerization of FGFR2. FLIM analysis of the FRET between the FGFR2-CFP and FGFR2-RFP. First panel: Reference lifetime measurements between FGFR2-GFP and RFP-alone, control for no interaction. The mean lifetime is centred around ∽2.1 ns (Blue), which corresponds to the mean lifetime for isolated CFP alone. Second panel: Dimerization of C1 showing a measurable left shift with of the molecule showing FRET above the control. Note that most interactions are seen in the intracellular vesicles. Third panel: Dimerization of C2. 16% of molecules on plasma membrane showing dimerization above the control threshold (Orange). Forth panel: Dimerization of C3, as with C1, 26% of the molecules are showing interaction (Orange) however unlike C1, almost all of the interactions are on the plasma membrane. Inserts with arrows showing exquisite separation of dimeric and non-dimeric FGFR2-C3 on the plasma membrane. (**D**) GST-CT^C1^ was used to pull down five mono-phosphorylated constructs of JM-KD^pY1^ with progressively truncated JM (JM-KD^pY1^, JM^Δ15^-KD^pY1^, JM^Δ25^-KD^pY1^, JM^Δ35^-KD^pY1^, and KD^pY1^). The presence of the intact JM enhances the interaction with GST-CT^C1^. (**E**) GST-CT^C2^ was used to pull down five mono-phosphorylated constructs of JM-KD^pY1^ as described in Fig. 6D. The presence of the intact JM also enhances the interaction with the CT from C2 isoform. (**F**) A GST-CT^C1^ pulldown of different FGFR2IIIb mono-phosphorylated proteins that include the presence or absence of JM and/or CT (KD^pY1^; JM-KD^pY1^; KD^pY1^-CT, and JM-KD^pY1^-CT), shows that the presence of JM, but not CT, promotes the interaction between kinase domain and GST-CT^C1^. The presence of CT inhibits the GST-CT^C1^ interaction, indicating CT binds through an intramolecular interaction. (**G**) A GST-JM pulldown of different FGFR2IIIb mono-phosphorylated proteins (as described in Fig. 5F), shows that the presence of JM does not block JM binding suggesting that JM of one protomer binds to the other in the mono-phosphorylated dimers (previously identified for KD and JM-KD). The latch to the protomer in the asymmetric dimer leaves an available JM binding site. The presence of CT (in KD-CT^C1^ and JM-KD^pY1^-CT^C1^) reduces JM binding. (**H**) MST measurement of JM binding to CT. A two-fold serial dilution of CT was titrated into JM which was labelled with Atto 488. (**I**) NMR titration of JM titrated into ^15^N-labelled CT using a red-to-white gradient, where white represents the weakest CSP and red depicts the strongest CSP. Proline residues are not visible in this experiment (shown in grey).

To confirm the impact of the CT on dimer formation in cells we used fluorescence lifetime imaging microscopy (FLIM) in serum starved HEK293T cells co-expressing CFP- and RFP-tagged FGFR2^C1^, FGFR2^C2^ and FGFR2^C3^. Compared to the control (FGFR2^C1^ with RFP) we saw increasingly shorter lifetimes in the populations of C2 and C3 receptors respectively, indicating that, in the absence of growth factor, dimerization increases in response to reduction in the size of CT (Fig. 5C). Interestingly the C3 isoform appears to be extensively membrane localized suggesting that recycling of the receptor was impaired (zoomed inset panels Fig. 5C). Together our data indicate that the binding of CT to active intermediate KD^pY1^ counters the positive impact of JM on RTK phosphorylation in the absence of stimuli by simultaneously binding to regions of KD inhibiting enzyme activity as well as restricting formation of the asymmetric dimer, i.e. involving both mechanisms hypothesised earlier.

### CT binds independently to JM

We have shown that CT binds more tightly in a 1:1 complex to JM-KD^pY1^ than to KD^pY1^ alone (Fig. 3C). We have also demonstrated that, although JM acts to sustain the asymmetric dimer and hence enhance activity, only the N-terminal residues of JM are involved in the intermolecular latch interaction. Thus, CT might be capable of binding to both KD^pY1^ and JM. To investigate a direct intramolecular interaction between JM and CT we first showed that binding of JM-KD^pY1^ to both CT^C1^ and CT^C2^ was reduced as JM was truncated (Fig. 5D and 5E respectively). Since stoichiometry of the final complex formed in each case is 1:1, the self-associated state of the JM-KD^pY1^ interactant should not produce the observed changes in binding. Interaction with CT was much reduced on deletion of residues 429 to 449 which are outside the region previously shown to bind to KD. Thus, it would be possible under certain conditions (e.g. within the context of the ligand bound full length receptor) that JM could maintain the latch interaction whilst simultaneously binding to CT. This suggests that CT has two independent modes of binding involved in RTK regulation: 1) binding to KD^pY1^ and inhibiting activity and dimerization, and 2) binding to JM without affecting latch formation. These modes will have distinct function, are mutually exclusive and occur at different time points in receptor up-regulation.

Using four mono-phosphorylated constructs; KD^pY1^, JM-KD^pY1^, KD^pY1^-CT^C1^, and JM-KD^pY1^-CT^C1^ in a pull-down assay with GST-CT^C1^, we showed that CT^C1^ binds independently to JM-KD^pY1^. However, when CT was included as part of the construct in both KD^pY1^-CT^C1^ and JM-KD^pY1^-CT^C1^ binding was abrogated (Fig. 5F). Thus, our data demonstrate that CT binds to JM-KD^pY1^ through an intramolecular interaction, since including CT on the construct disrupts dimerization and blocks GST-CT binding. We know that CT forms an extensive intramolecular interface with KD^pY1^ which stabilizes the monomeric protomer. This interface includes residues across the extent of the entire tail region (Fig. 3D). We have also shown that the presence of JM enhances the interaction with CT. Incubation of the same KD^pY1^, JM-KD^pY1^, KD^pY1^-CT^C1^, and JM-KD^pY1^-CT^C1^ with GST-JM showed that, consistent with previous observations, JM was able to bind to KD^pY1^ (Fig. 5G). JM also bound JM-KD^pY1,^ which, although dimerized through one JM latch, has a free KD for independent JM binding. Significant binding of JM to KD^pY1^-CT, but negligible binding of JM with the JM-KD^pY1^-CT^C1^ construct was observed. These interactions of JM in the presence of CT could not occur if CT successfully competed with JM for binding to KD, but would require that JM can bind simultaneously with KD and CT. We measured direct binding between JM and CT (K_d_ = 20.2 ± 2.92 µM; table S1 and Fig. 5H). We also identified that the highest affinity sequence of JM that recognized CT includes residues ^429^VTVSAESSSSMNSN^442^ (fig. S6A and table S1). Thus, the binding site on JM for CT is non-overlapping and C-terminal to the consensus sequence of JM that we showed is required for forming the intermolecular latch to KD^pY1^, i.e. ^414^PAVHKLT^420^, however it includes the VT site (including residues V429 and T430; the VT motif) (*34*) for FRS2 recruitment on growth factor binding. These data rationalize our previous cell-based observation that the presence of CT prevents FRS2 phosphorylation by FGFR2 under stimulated conditions (Fig. 3A).

To map the interaction between JM onto CT^C1^ we titrated unlabelled JM into ^15^N-labelled CT^C1^. The binding site can be seen to incorporate residues between V798 and S819 of CT which contains the proline-rich motif (Fig. 5I). Using a series of short peptides derived from CT we demonstrated that the proline-rich sequence from CT binds to JM (table S1 and fig. 6b), and the ^808^PCLPQYPH^815^ sequence is necessary for CT to bind to JM. Importantly, this is the same sequence that binds to both KD (Fig. 3) and to the GRB2 CSH3 domain (*35*). Thus, CT mediates three modes of receptor regulation.

**Fig. 6.**
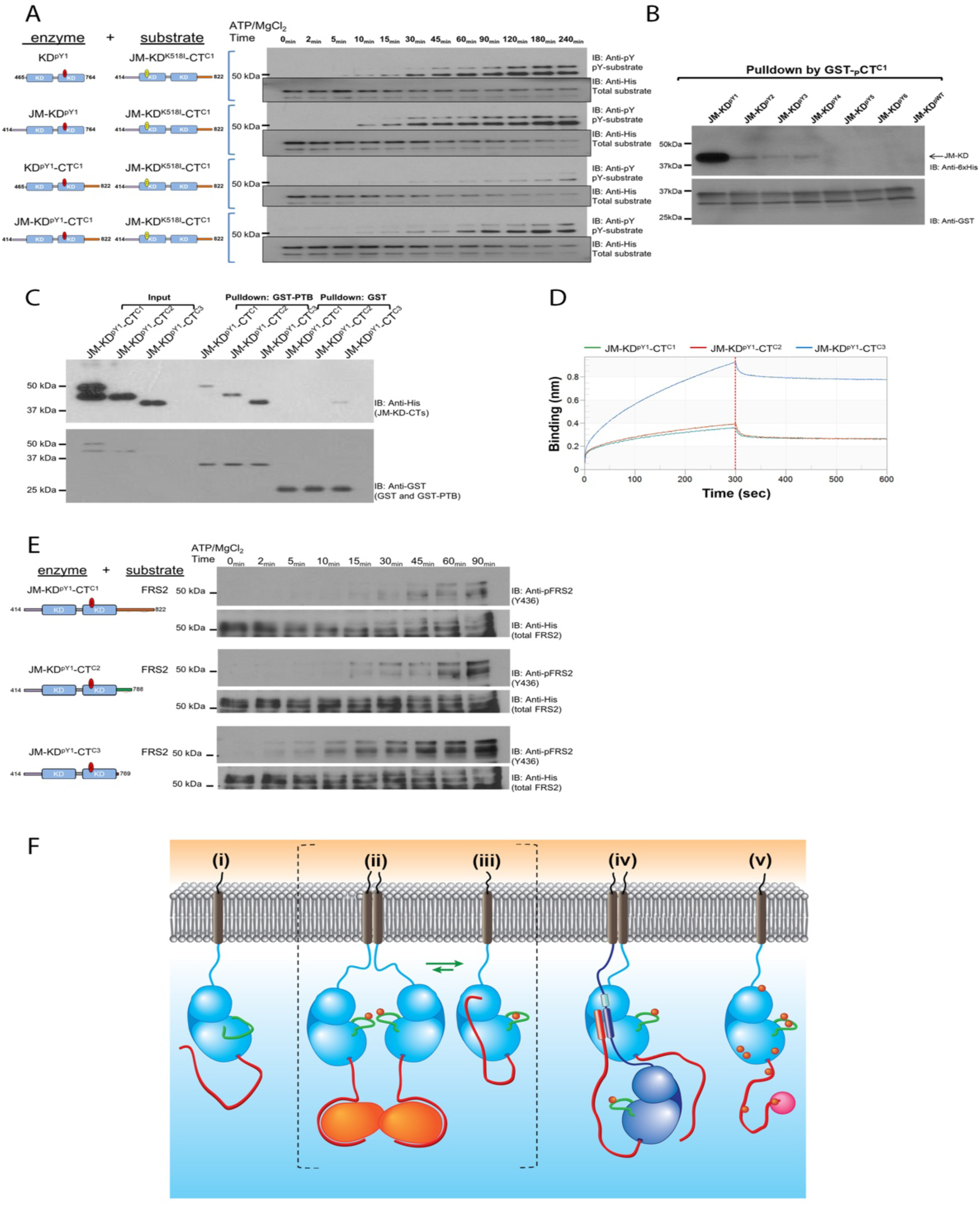
JM and CT combine to regulate KD. (**A**) Kinase activity is controlled by both JM and CT. KD^pY1^, JM-KD^pY1^, KD^pY1^-CT^C1^, and JM-KD^pY1^-CT^C1^, were incubated with kinase-dead JM-KD^K518I^-CT^C1^ in a 1:1000 ratio in the presence of ATP/Mg^2+^ and quenched with EDTA at different time points as indicated. The phosphorylation of JM-KD^K518I^-CT^C1^ was measured using a pY99 antibody. (**B**) Phosphorylation of CT (pCT^C1^) reduces its binding to KD with higher phosphorylation order. GST-CT^C1^ was phosphorylated by JM-KD^WT^-CT^C1^ and used for a GST pulldown assay with different phosphorylated JM-KD (JM-KD^pY1^ – JM-KD^pY6^ and wild type JM-KD). (**C**) GST-FRS2 PTB domain was used precipitate the following His-tagged constructs representing the mono-phosphorylated isoforms of FGFR2IIIb; JM-KD^pY1^-CT^C1^, JM-KD^pY1^-CT^C2^and JM-KD^pY1^-CT^C3^. The presence of the intact CT in the C1 isoform inhibits the interaction of FRS2 with its cognate site on JM. (**D**) BLI measurement of GST-FRS2 PTB binding to JM-KD^pY1^-CT^C1^, JM-KD^pY1^-CT^C2^ and JM-KD^pY1^-CT^C3^. The GST-PTB domain from FRS2 was immobilised on the sensor and was exposed to 2.6 µM of the cytoplasmic region of each of the FGFR2 isoforms. After 300 s the chip was washed. The sensorgrams clearly show that over the time course up to 300 seconds (prior to the washing step; dotted line), in the absence of CT (C3 isoform) a significantly increased amount of FGFR2 protein binds to the PTB domain compared with the C1 and C2 isoforms. (**E**) The FGFR2 isoforms (JM-KD^pY1^-CT^C1^, JM-KD^pY1^-CT^C2^ and JM-KD^pY1^-CT^C3^) were incubated with FRS2 protein in a 1:100 ratio in the presence of ATP/Mg^2+^ and quenched at different time points as indicated. The phosphorylation of FRS2 was measured using an anti-pFRS2 (Y436) antibody. (**F**) i: In the absence of stimulation the unphosphorylated FGFR2 (light blue, JM light blue line, CT red line) can exist as a monomer freely diffusing through the plasma membrane. ii: Random collision of FGFR2 results in dimer formation. Dimeric GRB2 (orange) is recruited via a proline-rich sequence on CT into a heterotetramer. This stabilizes the mono-phosphorylated active A-loop (green line) tyrosine residues (red circles) on KD, but signalling is stalled by the presence of GRB2 on CT. iii: the mono-phosphorylated KD provides a strong binding site for CT. CT to KD^pY1^ interaction results in the releasing of GRB2. This interaction inhibits the JM-mediated formation of asymmetric dimer and thus prevents the KD^pY1^ intermediate state from further auto-phosphorylation activity. Active intermediate states ii: and iii: are in equilibrium, the presence of the states is dependent on GRB2 concentration and hence the ability of GRB2 to compete with the intramolecular interaction with KD for binding to CT. iv: Binding of extracellular growth factor co-localizes two receptors and permits the formation the active, asymmetric dimeric conformation. This is sustained by the interaction of JM from the enzyme-like receptor (dark blue) with KD of the substrate-like receptor (light blue). The sequence on JM which binds to KD (light blue thick line) is immediately proximal to a sequence (dark blue thick line) which binds in an independent interaction to CT of the enzyme-like receptor. This binding site (red thick line) includes the proline-rich motif that recognizes a site on KD and GRB2. Thus, JM-CT interaction blocks auto-inhibition and GRB2 recruitment. This ensures that the active state is prolonged. In this conformation the CT of the substrate-like receptor can access the enzyme-like KD active site. v: Prolonged activity of the dynamic asymmetric dimer results in increasing phosphorylation of KD and CT. As the pY burden increases the dimerization between KDs reduces until they fully dissociate. The phosphorylated KD abrogates the inhibitory intramolecular binding of CT and the recruitment of GRB2. The receptor is therefore available for recruitment of downstream effector proteins (magenta).

The ability of CT to bind mutually exclusively to both KD and JM suggests that it can adopt two distinct conformations which have opposing impact on kinase activity, and which operate independently at different time points in the receptor up-regulation process. 1) Binding of CT to KD^pY1^ results in an intramolecular auto-inhibitory conformation which sustains the monomeric state of the mono-phosphorylated receptor. This is expected to control the active intermediate state in the absence of growth factor stimulation. 2) The CT of the enzyme-like KD flips from the inhibitory interaction with KD^pY1^, to bind intramolecularly to the JM. In this state JM can simultaneously perform the role as the latch to the substrate-like KD of the asymmetric dimer. Thus, the CT is effectively isolated from blocking the kinase activity and receptor dissociation. Also in this state the CT of the substrate-like protomer is free to become phosphorylated. We propose that CT is stabilized in the latter of these two conformations when the receptor is bound to extracellular growth factor. The direct interaction between two peripheral regions thus adds a level of control to the kinase output not previously observed for RTKs.

### KD can exist in equilibrium between Grb2-bound and CT-bound active intermediate states

Previously we have shown that in the absence of growth factor FGFR2 binds to the C-terminal SH3 domain of dimeric GRB2 through the proline-rich sequence on CT (^808^PCLPQYP^814^) and maintains FGFR2 in an active intermediate, signalling incompetent heterotetrameric state (*4, 36*). Since we have shown that the proline-rich sequence binds intramolecularly to JM-KD^pY1^, the ability of CT to regulate the active intermediate state is manifold. To assess whether these interactions are mutually exclusive, we first performed a pulldown assay using GST-CT which was incubated with GRB2. Subsequent addition of increasing concentrations of JM-KD^pY1^ gradually decreased the amount of GRB2 precipitated, suggesting JM-KD^pY1^ competes with GRB2 for the binding site on CT (fig. S6C). Based on the respective measured affinities, it is assumed that CT was binding to KD rather than JM in this experiment. CT was then immobilized on biolayer interferometry (BLI) sensors to probe the competitive interactions of JM-KD^pY1^ and GRB2. Binding of JM-KD^pY1^ or GRB2 alone to the immobilized CT showed the expected different interaction properties associated with the size of the added proteins (fig. S6D) however, the addition of both JM-KD^pY1^ and GRB2 simultaneously gives an identical response to that of adding only JM-KD^pY1^ to immobilized CT (fig. S6D), suggesting that GRB2 competes less efficiently with JM-KD^pY1^. Sequential binding of JM-KD^pY1^ and GRB2 to the immobilized CT (fig. S6E) also indicated that, once CT is saturated with JM-KD^pY1^, GRB2 cannot compete it off. Interestingly, application of JM-KD^pY1^ to GRB2-saturated CT sensor resulted in a decrease of the binding signal followed by an increase of interaction signal as the JM-KD^pY1^ displaces GRB2 on the immobilized CT (fig. S6F). Thus, equilibrium between CT bound to GRB2 and CT bound to JM-KD^pY1^ exists but the respective affinities and the intramolecular nature of interaction strongly favours the latter.

### JM and CT regions combine to regulate kinase activity

Having demonstrated the positive and negative impacts of JM and CT binding on regulation of KD^pY1^ respectively, we examined how the presence of these peripheral regions control FGFR2 signalling. The following constructs, KD^pY1^; JM-KD^pY1^; KD^pY1^-CT^C1^; and JM-KD^pY1^-CT^C1^ were used in an assay with the kinase dead JM-KD^K518I^-CT as a substrate (Fig. 6A). The activity is slightly enhanced by the presence of JM in JM-KD^pY1^ compared to KD^pY1^. This is consistent with JM stabilizing the active asymmetric dimer. Conversely, the presence of CT in KD^pY1^-CT^C1^ dramatically inhibits kinase activity through previously observed direct interaction with KD and resulting inhibition of dimerization. The presence of JM and CT in JM-KD^pY1^-CT^C1^shows medium activity. This underscores the regulatory role of the interplay between the two peripheral regions of the receptor in sustaining the active intermediate state through modulation of kinase activity. These data mirror the experiments on the constitutively active truncated K-*sam* FGFR2 isoforms, where the absence of CT leads to dysregulation of the intact receptor (Fig. 3A).

As the receptor becomes progressively phosphorylated the interaction of CT to KD needs to be down-regulated to enable access of downstream signalling proteins. We used a pull down experiment to reveal the mechanism for this release. GST-CT which was phosphorylated on its available tyrosine residues, pCT^C1^, was unable to pulldown JM-KD as it became progressively phosphorylated, i.e. JM-KD^pY1^ to JM-KD^pY6^ (Fig. 6B) Thus, as the receptor pY load increases, the phosphorylated CT is less able to bind intramolecularly, making it available for recruitment of downstream effector proteins.

We have shown that residues in JM sequence 429 to 449 play a role in interacting with CT (Fig. 5D), and hence the presence of CT would occlude the ^429^VT^430^ binding motif (*34*) for the FRS2 phosphotyrosine binding domain (PTB) in the absence of stimulation and prevent aberrant signalling. Therefore, the absence of CT in the C3 isoform should allow unrestricted recruitment of FRS2 leading to prolonged phosphorylation even at the unliganded state (as seen in Fig. 3A). This inhibition of access of FRS2 to JM by CT was demonstrated where significantly less JM-KD^pY1^-CT^C1^ was precipitated compared to JM-KD^pY1^-CT^C3^ in a pull down assay using GST-FRS2 PTB domain (Fig. 6C). JM-KD^pY1^-CT^C2^ showed an intermediate level of interaction consistent with the proline-rich motif present in this isoform binds with lower affinity to the FRS2 cognate site. We also measured different FGFR2 isomers binding to FRS2 using BLI. In the absence of the intact CT (C3 isoform) a significantly increased amount of FGFR2 protein bound to the PTB domain compared with the C1 and C2 isoforms (Fig. 6D).

Using an *in vitro* kinase assay we were able to demonstrate that the phosphorylation of FRS2 by FGFR2 is affected by CT in the different isoforms. Immunoblotting showed that the C3 isoform has the highest kinase activity toward FRS2, whereas the C1 isoform has the lowest (Fig. 6E). This further suggested that CT^C1^ isoform can interact with JM which contains the ^429^VT^430^ motif, and inhibit the recruitment and phosphorylation of FRS2 in the active intermediate state. This observation explains why the C3 isoform has higher FRS2-mediated downstream signalling activity and exhibits uncontrolled activation leading to oncogenic outcome in the active intermediate state.

### General importance of proline-rich motifs in RTK regulation

Having shown that proline-rich motifs are critical in regulatory interactions with JM and KD, and the mutations on proline residues affect FGFR2 kinase activity and downstream signalling, we investigated whether mutation/truncation of proline-rich motifs within CTs of other RTKs are found in cancers in general. Humans have 58 identified RTKs, which fall into twenty subfamilies (*10*) of which 49 have proline residues on their CTs. Apart from providing protein recruitment sites for SH3/WW domains, the importance of proline-rich motifs in RTK signalling has not been investigated. Genomic data from cancer patient samples available on cBioPortal for Cancer Genomics (www.cbioportal.org) shows that of the 49 RTKs, 40 have been identified with proline residue point mutations or deletions of proline-containing CT sequences (table S4). This suggests an important role for proline-containing sequences and raises their importance in regulation, particularly during the non-stimulated, active intermediate state we present here.

### Discussion

RTKs generally cycle through a series of states on going from the dephosphorylated monomeric state to the fully phosphorylated, signalling-competent state. We have investigated a series of snapshots of structural states which show possible interactions and juxtapositioning of the various components. Linking of these snapshots into an animated series permits a full understanding of the progression of events and how each one provokes the next. Our data have highlighted RTKs in the mono-phosphorylated active intermediate JM-KD^pY1^-CT state as the most important frame in the animation of the progress from inactive to signalling receptor. This state represents a major check-point because the KD is active, but signal transduction is inhibited. Our observations show how, in this state, the receptor is highly regulated by the interactions of JM and the proline-rich motif on CT in the absence of stimulation, whilst being primed for full activation on growth factor binding.

Our mechanistic model starts with the dephosphorylated, inactive monomeric state, and progresses via the mono-phosphorylated active intermediate, through to the fully active state (Fig. 6F). Under basal conditions growth factor receptors diffuse through the plasma membrane (Fig. 6F i). Random collision leads to transmembrane-mediated self-association. This can lead to as much as 20% dimer and subsequent A-loop tyrosine phosphorylation of FGFR2 in the absence of ligand (*3*). Binding of GRB2 stabilizes FGFR2 dimers in the active intermediate state and inhibits further phosphorylation (*4*) (Fig. 6F ii). The presence of GRB2 on CT of FGFR2 is perturbed by a cycle of phosphorylation by the receptor and subsequent dephosphorylation by SHP2 phosphatase (*4, 36*). Without the inhibitory function of GRB2, the dysregulated active intermediate receptor could progress to full activation through the formation of asymmetric dimer mediated by JM. This would leave receptor activation in a precarious position without an additional mechanism restrict further kinase activity. Our observations demonstrate that this mechanism involves the intramolecular binding of CT to JM-KD^pY1^ (Fig. 3C). This interaction is mutually exclusive of binding of GRB2 (fig. S6C to S6F and Fig. 6F iii). The presence of CT inhibits further phosphorylation by inhibiting catalytic activity and asymmetric dimerization. Thus, the mono-phosphorylated state, which is the check-point prior to full receptor activation, is tightly regulated either by the binding of GRB2 or the interaction with CT.

When cells are exposed to extracellular stimulation, JM of one protomer in the dimer latches onto KD of the other (Fig. 6f iv). In this way the former becomes the designated enzyme-like receptor, whilst the latter becomes the substrate-like receptor, both being held in a moderate affinity, dimeric conformation. Since the KD is already in its mono-phosphorylated state it is available for JM binding and adoption of the asymmetric dimer conformation. JM appears to inhibit the direct interaction between KDs and promote dynamic interlocution between the active domains in an asymmetric dimer (Fig. 2C). It also juxtaposes KDs within the asymmetric dimer to prevent progressive oligomerization of the domains (i.e. ‘daisy chain’ formation (*18, 37*)) allowing only dimers to form.

The unrestricted activity of the enzyme-like receptor requires that CT is not able to bind to, and hence down-regulate KD of this protomer. This state is achieved through the intramolecular binding of CT to the available site on JM. Thus, the role of CT is multi-faceted and functions through independent interactions with GRB2, KD and JM. Truncation of CT up-regulates the kinase (Fig. 3A and 3B) and hence provides a rationale for the elevated proliferative signalling as we seen in the oncogenic K*sam* deletions (fig. S6G).

The release of negative control by CT promotes an increase in KD phosphorylation which weakens dimerization (Fig. 2A), enhancing the dynamic interplay between protomers permitting easier access to tyrosine sites and alternation of the enzyme-like and substrate-like states between the molecules. Increasing phosphorylation of KD also results in progressive weakening of interactions with the peripheral regions ultimately leading to kinase dimer dissociation, leaving the receptor in a highly phosphorylated state whereby it can recruit downstream effector proteins (Fig. 6F v). Dissociation of phosphorylated FGFR2 also leaves it exposed to phosphatase activity which ultimately returns it to its initial unphosphorylated state (Fig. 6F i). Clearly the controlled activation cycle of FGFR2 would be affected by the impact of additional factors such as endocytosis (*33*), fluctuations in GRB2 concentration (*38*) and phosphatase concentration (*36*). However, the importance of both peripheral regions in influencing the self-association, and the dimeric conformation underscores how the receptor is tightly regulated to avoid aberrant signalling during the progression from unphosphorylated inactive form to fully phosphorylated active form.

Different RTKs include features that enable idiosyncratic regulation and commitment to defined downstream outcomes. The presence of peripheral regions on the majority of RTKs suggests that these can play common roles. In particular, the prevalence of proline-containing sequences in CTs. The impact on kinase-driven pathology of their mutation/truncation highlights the regulatory importance of these sequences. Mutations within, or deletions of proline-rich sequences in CTs of many RTKs (including IGF1R, FGFR2, ERBB2, ERBB3, ERBB4 and ROR2) are associated with range of cancers (*27*–*29*) (table S4). We hypothesise therefore that, since these sequences are found in the majority of RTKs, their interaction with KD and/or JM is a common feature of RTK regulation. Our data show, in the case of the K-*sam*II truncations, that when these regulatory features are perturbed pathogenicity can result in uncontrolled cellular signalling. Thus, understanding of the roles of the peripheral region interactions will suggest alternative routes for therapeutic intervention outside the currently well-trodden path of inhibition of kinase activity.

## Materials and Methods

### Cell culture

HEK293T cells were maintained in DMEM (Dulbecco’s modified Eagle’s high glucose medium) supplemented with 10% (v/v) FBS (foetal bovine serum) and 1% antibiotic/antimycotic (Lonza) in a humidified incubator with 10% CO_2_.

### Protein expression and purification

All MBP-tagged, GST-tagged and 6xHistidine-tagged fusion proteins were expressed and purified from BL21(DE3) cells. A single colony was used to inoculate 100 mL of LB which was grown overnight at 37°C. 1L of LB was inoculated with 10 mL of the overnight culture and allowed to grow at 37°C until the OD_600_ reaches 0.8 at which point the culture was cooled down to 20°C. Expression was then induced with 0.5 mM IPTG and the culture was grown for a further 12 hours before harvesting by centrifugation. Cells were re-suspended in 20 mM Tris, 150 mM NaCl, 10% glycerol, pH 8.0 in the presence of protease inhibitors and lysed by sonication. Insoluble material was removed by centrifugation (40,000g at 4°C for 60 min). The soluble fraction was applied to an appropriate affinity column (Amylose column for MBP-tagged proteins, GST column for GST-tagged proteins and Talon column for His-tagged proteins). Following a wash with 10 times column volume of wash buffer (20 mM Tris, 150 mM NaCl, pH 8.0), the protein was eluted from the column with elution buffer (the washing buffer supplemented with 20mM maltose for the MBP-tagged proteins; a supplement of 20mM reduced glutathione for the GST-tagged proteins; a supplement of 150mM imidazole for the 6xHis-tagged proteins) and was concentrated to 5 mL and applied to a Superdex 75 gel filtration column equilibrated in a buffer containing 20 mM HEPES, 150 mM NaCl and 1 mM TCEP pH 7.5. Analysis for protein purities by SDS-PAGE showed greater than 98% purity. For CT (CT^C1^ and CT^C1Δ34^, GST-tagged) production and JM-KD^pY1^-CT^C1^ (for crystallography, 6xHis-tagged), 1 unit of thrombin (Sigma T6884) was used to cleave 1mg of recombinant proteins at 4°C for overnight. After cleavage, Benzamidine Sepharose 4 Fast Flow beads (GE) were used to remove thrombin. GST-Tag/His-Tag and uncut proteins were removed by passing protein solution through a GST or Talon column. Expression of ^15^N-labelled proteins for NMR titrations and ^2^H, ^15^N, ^13^C-labelled protein for backbone resonance assignment was done as previously described(*39*). For expression in 100% D_2_O, this procedure was modified by pre-growing the culture in a small volume of 100% D_2_O prior to expression over 20 hours.

### Nuclear magnetic resonance (NMR) spectroscopy

#### General information

All NMR spectroscopic experiments concerning KD backbone assignment were carried out on Bruker Avance III 950 MHz NMR spectrometers equipped with cryogenically cooled triple resonance probes (5mm TXO or 3mm TCI). Titration experiments were additionally carried on Bruker Avance III 750 MHz NMR spectrometer, equipped with ^1^H-optimized triple resonance NMR 5mm TCI-cryoprobe. NMR data was processed using NMRPipe and further analyzed with CcpNmr Analysis software package available locally and on NMRBox platform. Chemical shift perturbations (CSPs) were calculated from the chemical shifts of backbone amide ^1^H (Δω_H_) and ^15^N (Δω_N_) using the following equation: 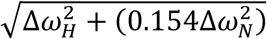

#### KD and CT backbone assignment

In order to obtain backbone assignment of KD and CT, both proteins were expressed in isotopically labeled media as described above. Set of KD samples uniformly ^13^C-^15^N labeled and fully or partially (70%) deuteriated (^2^H) as well as uniformly ^13^C-^15^N-labeled CT were prepared. Spectra were recorded at 25°C with KD concentrations ranging from 200 μM to 600 μM in a HEPES buffer (20 mM HEPES, 150 mM NaCl and 1 mM TCEP pH 7.5). Spectra of CT were recorded using 300 μM sample in the same HEPES buffer. Standard Bruker library together with BEST versions of amide transverse relaxation optimized spectroscopy (TROSY) of 3D backbone resonance assignment pulse sequences (HNCA, HNCOCA, HNCACB, CACBCONH, HNCO and HNCACO) were applied to collect high resolution spectra. In order to shorten acquisition time, Non-Uniform Sampling (20-30%) was routinely used.

#### KD titration with CT and CT-derived fragments

The NMR titration of uniformly ^15^N-labeled KD with unlabelled CT and CT-derived peptides were recorded at 25°C using 100 μM KD sample in HEPES buffer. CT and CT-derived fragments were added at the 1:0.5, 1:1, 1:1.5, 1:2 and 1:1, 1:5, 1:10 molar ratios respectively and the amide spectra were recorded using BEST TROSY pulse sequence.

#### CT titration with preform KD-JM complex

The NMR titration of KD with JM peptide were recorded at 25°C using 250 μM uniformly ^15^N-labeled sample in HEPES buffer. JM peptide was added at the 1:1, 1:2 and 1:3 molar ratios and the amide spectra recorded using BEST TROSY pulse sequence. To the fully titrated KD, peptide JM was added and the amide spectra (BEST TROSY) recorded at 1:1, 1:2, 1:3 molar ratios.

#### CT titration with KD

The ^15^N-labeled CT sample concentrated to 300 μM in HEPES buffer was titrated with unlabelled KD. Amide spectra were recorded at 25°C using hsqcetfpf3gpsi pulse sequence from Brüker library at 2:1, 1:1, 1:2, 1:3, 1:4, 1:8 and 1:12 molar ratios.

### X-ray crystallography

Crystals of JM-KD^pY1^-CT^C1^ were obtained using the hanging-drop vapour diffusion method, mixing equal volumes of protein with reservoir solution and equilibrating over this reservoir at 20°C for 2 weeks. The reservoir solution contained 100 mM Tris, 160 mM TMAO, 20% PEG2000 at pH 8.6. For cryoprotection, crystals were transferred in the crystallization buffer supplemented by 20% Glycerol. X-ray diffraction data sets were collected from frozen single crystals at the Advanced Light Source (Berkley, CA, USA, beamline 8.3.1) and processed with the program Elves. A molecular replacement solution was obtained using the BALBES molecular replacement pipeline and the crystal structure PDB code 2PSQ. Iterative model rebuilding and refinement was performed by using the program COOT, REFMAC5 and PDB_REDO against the data set. Structural figures were made using PyMol.

### Small-Angle X-ray Scattering (SAXS)

Data were collected for the mono-phosphorylated forms of KD^pY1^ and JM-KD^pY1^ on the SIBYLS beamline at the Advanced Light Source, Berkeley, USA at a wavelength of 1 Å. Every sample was exposed successively for 0.5, 1.0, and 6.0 s. Protein sample concentrations ranged from 1.0–10 mg/mL. Data for protein sample and buffer alone and were recorded at 10°C, and the buffer contribution was subtracted from the protein scattering data. Additional exploratory SAXS data were recorded at the SWING beamline (SOLEIL, Saint-Aubin, France) at λ = 1.03 Å. Data were analysed using PRIMUS, GASBOR, DAMMIF, and DAMAVER from the ATSAS software package 1. Model SAXS patterns were calculated and fitted to data using FoXS 2. MultiFOXS was used to evaluate mixtures of monomers and dimers.

Swiss-Model was used to prepare a complete FGFR2 KD^pY1^ monomer (including loops and side chains missing in the crystallographic model) based on PDB 2PVF. Potential FGFR2 KD^pY1^ dimers were based on arranging the completed monomer into dimer configurations found in crystal structures: PDB 2PSQ (symmetric head-tail dimer); 2PVF and 3CLY (asymmetric kinase C-terminal tail trans-phosphorylation); 3GQI (asymmetric A-loop trans-phosphorylation of FGFR1). Additional dimers were derived from our FGFR2 KD^pY1^ crystal structures (PDB 6V6Q). These crystals were formed by a combination of the symmetric head-tail dimers found in 2PSQ, as well as two other forms. In one, termed KI, contacts were established through the kinase insertion (KI) loop. In the other, termed N-C, the N-lobe of one kinase bound to the C-lobe of the other (See Supplementary Fig. 2d). 30 additional dimers were obtained using in silico docking with CLusPro2.0 (30 different docking results) based on monomeric KD. An ‘aggregate’ was mimicked using a region of the crystal lattice that had a similar structure as DAMMIF ab initio models derived from SAXS data under conditions where KD was highly aggregated.

We arranged these models into three different ‘pools’ of structures from which best fitting multi-state models were subsequently computationally selected (using FoXS). Pool 1: all monomeric and dimeric models (44 models). Pool 2: monomer and only dimers found in our FGFR2^pY1^ crystal structures (7 models). Pool 3: As Pool 2, but also including trimers derived from our FGFR2^pY1^ crystal lattices (9 models). For JM-KD^pY1^ We prepared models similarly as for KD^pY1^, using crystal-derived and in silico docked dimeric and multimeric assemblies. Given that the long flexible JM region has a significant impact on the SAXS pattern, we typically used five different JM conformations for the monomer and crystal-lattice derived dimers, resulting in a pool of 55 different structures.

### Mutation of FGFR2 proteins

Standard site-directed mutagenesis was carried out to mutate tyrosine residues into phenylalanine on KDs to mimic the sequential phosphorylation pattern of KD (KD^pY1^ to KD^pY6^; see schematic Fig. 1D and Supplementary Fig. 2A. For the FGFR2 IIIb isoform this sequence is; pY657, pY587, pY467, pY589, pY658 and pY734, adapted from the IIIc isoform(30)). The same methods were also used to mutate proline residues on the C-terminal tail in this study.

### In vitro dephosphorylation and phosphorylation of purified proteins

Calf Intestinal alkaline phosphatase (CIP, New England Biolabs) was conjugated on UltraLink Biosupport beads (Thermo Fisher Scientific). CIP-beads were mixed with purified protein solution and rotated gently at 4°C for overnight to remove phosphate group in solution. After dephosphorylation, protein solution and CIP-beads were separated by centrifugation. Dephosphorylation level was examined by western blotting. Purified FGFR2 proteins were phosphorylated by incubating with 5 mM ATP and 10 mM MgCl2. The phosphorylation reactions were quenched by adding EDTA (prepared in 10 mM HEPES, pH 7.5) to a final concentration of 100 mM. Proteins were analysed by SDS-PAGE and western blot to study the phosphorylation status.

### Transient cell transfection with plasmids

30 min before transfection, cells were harvested and resuspended in antibiotic-free medium. Transfection was carried out using Metafectene (Biontex Cat#: T020) according to manufacturer manual.

### Cell signalling studies

For mammalian cell studies, cells were starved for 16 hours, and left unstimulated or stimulated with 10ng/ml FGF7 ligand (R&D Systems Cat#: 251-KG/CF) at 37°C. After stimulation, medium was removed and cells were put on ice and immediately lysed by scraping in ice-cold lysis buffer supplemented with protease inhibitor (Calbiochem) and phosphatase inhibitor (1 mM sodium orthovanadate (NaVO3), and 10 mM sodium fluoride (NaF). Cells were cleared by centrifugation and the supernatants were subjected to immunoblotting using the BioRad protein electrophoresis system. The intact gel was transfer to PVDF membrane for probing with different antibodies. Phospho-protein blots were stripped with stripping buffer (Millipore) and re-probed with total protein antibodies. Antibodies were from: Anti-Phospho-FGF Receptor (Tyr653/654) Rabbit polyclonal, Cell Signaling Technology Cat#:3471; Anti-Phospho-FRS2-α (Tyr436) Rabbit polyclonal, Cell Signaling Technology Cat#: 3861; Anti-Phospho-SHP-2 (Tyr542) Rabbit polyclonal, Cell Signaling Technology Cat#: 3751; Anti-Phospho-Akt (Thr308) Rabbit monoclonal, Cell Signaling Technology Cat#: 4056; Anti-Phospho-p44/42 MAPK (Erk1/2) (Thr202/Tyr204) Rabbit monoclonal, Cell Signaling Technology Cat#: 4370; Anti-p44/42 MAPK (Erk1/2) Rabbit monoclonal, Cell Signaling Technology Cat#: 4695; Anti-GRB2 Rabbit polyclonal, Cell Signaling Technology Cat#: 3972; Anti-α-Tubulin Rabbit polyclonal, Cell Signaling Technology Cat#: 2144; Anti-GST Rabbit polyclonal, Cell Signaling Technology Cat#: 2622; Anti-FGFR2 Mouse monoclonal, Santa Cruz Biotechnology Cat#: sc-6930; Anti-Phospho-Tyr Mouse monoclonal, Santa Cruz Biotechnology Cat#: sc-7020; Anti-6xHis Mouse monoclonal, Takara Cat#: 631212.

### Pulldown and western blots

For immunoblotting, proteins were separated by SDS-PAGE, transferred to PVDF membranes and incubated with the specific antibodies. Immune complexes were detected with horseradish peroxidase conjugated secondary antibodies and visualized by enhanced chemiluminescence reagent according to the manufacturer’s instructions (Pierce). For pulldown experiments, 100ug of protein was prepared in 1 ml volume. MBP-tagged or GST-tagged proteins immobilized on Amylose beads (GE Healthcare Life Science) or Glutathione Sepharose (GE Healthcare Life Science) was added and incubated at 4°C overnight with gentle rotation. The beads were then spun down at 4,000 rpm for 3 minutes, supernatant was removed and the beads were washed with 1 ml lysis buffer. This washing procedure was repeated five times in order to remove non-specific binding. After the last wash, 50 µl of 2x Laemmli sample buffer were added, the sample was boiled and subjected to SDS-PAGE and western blot assays.

### Fluorescence resonance energy transfer (FRET)

Recombinant GFP-JM-KD^pY1^-CT^C1^ (donor) and RFP-JM-KD^pY1^-CT^C1^ (acceptor) proteins (1µM) were used for In vitro steady-state FRET analysis. The changes of donor emission (510nm) upon dimer formation or dimer disruption upon the addition of CT were recorded at 25°C.

### Quantitative imaging FRET microscopy

HEK293T cells 24 h after transfection were seeded onto coverslips and allowed to grow for a further 48 h then fixed by addition of 4% (w/vol) paraformaldehyde, pH 8.0, 20 min. at room temperature. Cells were then washed six or seven times with PBS, pH 8.0 and mounted onto a slide with mounting medium (0.1% p-phenylenediamine/ 75% glycerol in PBS at pH 7.5 – 8.0) and curated for 3 - 4 h before imaging. FLIM images were captured using a Leica SP5 II confocal microscope. Atto488 was excited at 900 nm with titanium–sapphire pumped laser (Mai Tai BB, Spectral Physics) with 710 - 920nm tunability and 70 femtosecond pulse width. Becker & Hickl (B&H) SPC830 data and image acquisition card was used for time-correlated single photon counting (TCSPC). Electrical time resolution 8 Pico seconds with a pixel resolution of 512 x 512. Data processing and analysis were done using B&H SPC FLIM analysis software. The fluorescence decays were fitted with a single exponential decay model.

### Microscale thermophoresis (MST)

Binding affinities were measured using the Monolith NT.115 (NanoTemper Technologies, GmbH). Proteins were fluorescently labelled with Atto488 according to the manufacturer’s protocol. Labelling efficiency was determined to be 1:1 (protein:dye) by measuring the absorbance at 280 and 488 nm. A 16 step dilution series of the unlabelled binding partner was prepared and mixed with the labelled protein at 1:1 ratio and loaded into capillaries. Measurements were performed at 25 °C in a buffer containing 20 mM HEPES, 150 mM NaCl, 1 mM TCEP and 0.01% Tween 20 at pH7.5. Data analysis was performed using Nanotemper Analysis software, v.1.2.101 and was plotted using Origin 7.0. All measurements were conducted as triplicates and the error bars were presented as the standard deviations of the triplicates. For the experiments employed to measure dimerization KD values are referred to a ‘apparent’ because, based on the differential concentrations, the fitting model assumes labelled are bound to unlabelled polypeptides.

### Surface plasmon resonance (SPR)

SPR experiments were carried out using a BIAcore T100 instrument (GE Healthcare). CT^C1^ were immobilized on CM4 chips according to the standard amine coupling protocol. Briefly, carboxymethyl groups on the chip surface were activated with a 1:1 mixture of N-ethyl-N-(dimethyaminopropyl) carbodiimide (EDC) and N-hydroxysuccinimide (NHS). Proteins were diluted in 20 mM HEPES, pH 6.5 and injected over the activated chip surface. The unbound chip surface was blocked using ethanolamine. Proteins were immobilized to approximately 200 response units. Different concentrations of analytes were injected over the immobilized chips at a flow rate of 30 μl/min. The sensor surface was regenerated by injection of 30 μl of 0.1% SDS and 60 μl of 500 mM NaCl. Reference responses were subtracted from flow cells for each analyte injection using BiaEvaluation software. The resulting sensorgrams were anaylsed to determine the kinetic parameters. Raw data shows a rise in signal associated with binding followed by a diminished signal after application of wash buffer.

### Bio-layer interferometry (BLI)

BLI experiments were performed using a FortéBio Octet Red 384 using Anti-GST sensors. Assays were done in 384 well plates at 25 °C. Association was measured by dipping sensors into solutions of analyte protein (FGFR2 proteins) for 125 seconds and was followed by moving sensors to wash buffer for 100 seconds to monitor the dissociation process. Raw data shows a rise in signal associated with binding followed by a diminished signal after application of wash buffer.

### Peptides

^407^Juxtamembrane region^462, 407^KPDFSSQPAVHKLT^420, 414^PAVHKLTKRIPLRRQVT^430, 429^VTVSAESSSSMNSN^442, 439^MNSNTPLVRITTRL^452, 449^TTRLSSTADTPMLA^462, 801^PDPMPYEP^808^,^801^PDPMPYEPCLPQYPH^815, 808^PCLPQYPHINGSVKT^822, 801^PDPMPYEPCLPQYPH^815^,^808^PCLPQYPH^815, 808^PCLPQYPHINGS^819, 815^HINGSVKT^822, 804^MPYEPCLP^811^. All peptides were purchase from Genscript.

## Supplementary Materials

Fig. S1. The interaction of JM to KD.

Fig. S2. Dimerization of mono-phosphorylated kinase.

Fig. S3. CT^C1^ fragments bind to KD^pY1^.

Fig. S4. Binding of CT does not compete with JM for binding to KD^pY1^.

Fig. S5. Phosphorylation states control KD and CT^C1^ interaction.

Fig. S6. CT-JM interactions and CT competes with GRB2 for binding to JM-KD^pY1^.

Table S1. Biophysical measurements of FGFR2 JM, KD, and CT interactions.

Table S2. Statistic parameters of SAXS experiments for KD^pY1^ and JM-KD^pY1^.

Table S3. X-ray data collection and refinement statistics.

Table S4. Proline mutants in RTK C-terminal tails and human cancers.

## Acknowledgments

We thank the Berkeley Laboratory Advanced Light Source and SIBYLS beamline staff at 12.3.1 for assistance with collection of SAXS data, and we acknowledge SOLEIL for provision of synchrotron radiation facilities (proposals nr. 20181104 and 20190107) and we would like to thank J. Perez and A. Thureau for assistance in using the beamline SWING. We thank Dr S. Arur (MD Anderson Cancer Center), Dr N. Forde (University of Leeds), and A. Stainthorp (University of Leeds) for helpful discussion and comments. We thank Dr A. Kalverda (The Astbury Structural Biology Laboratory, University of Leeds) for the help on NMR data collection and analysis.

## Author contributions

C.-C.L. and J.E.L. designed the overall project and wrote the manuscript. C.-C.L. performed and analysed most of the experiments. L.W. assigned and analysed NMR data. K.M.S. contributed to data analysis and manuscript writing. S.T.A and G.P-M. performed the protein structural analysis and contributed to manuscript writing. Z.A. performed the FLIM experiments and data analysis.

## Data and materials availability

The accession number for the coordinate and structure for the mono-phosphorylated FGFR2 kinase reported in this paper is PDB: 6V6Q.

## Supplementary Materials

**Fig. S1.**
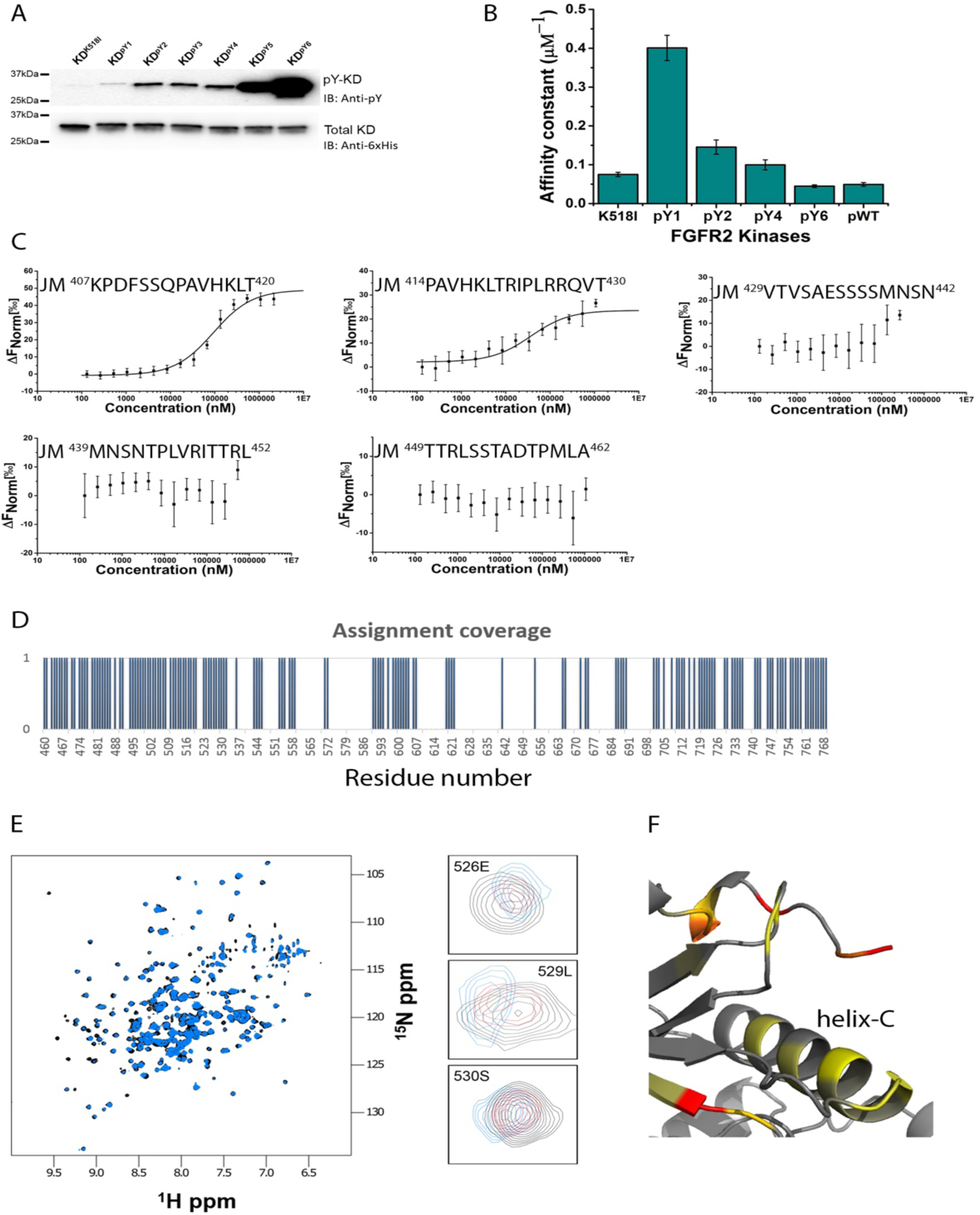
The interaction of JM to KD. (**A**) The phosphorylation states of KD^pY1^ to KD^pY6^ were confirmed using a phosphotyrosine pY99 antibody. An anti-6xHis tag antibody was used to probe for total proteins as the loading control. (**B**) Quantification of binding affinities between JM and KD with different phosphorylation states. The mono-phosphorylated KD is the strongest binding partner for JM. (**C**) Five short JM peptides were synthesised (residues 407-420, 414-430, 429-442, 439-452, and 449-462) and used to identify the binding region for KD^pY1^. The MST measurement results indicate that residue 407-420 provides the best binding ability for KD^pY1^. (**D**) Assignment coverage of KD amide backbones. Overall percentage of the assignment used in all titration experiment was 56% (not including prolines).(**E**) ^1^H-^15^N HSQC spectra of unbound mono-phosphorylated FGFR2 kinase KD^pY1^ (black) overlay with JM bound KD^pY1^ (blue) at 1:2 ratio. Examples of peaks with high chemical-shift perturbations (CSPs) are shown by labels indicating the assignment of given peaks. (**F**) CSPs of JM binding to KD^pY1^ plotted on the X-ray structure of mono-phosphorylated kinase. Minor CSPs appear on one side of the regulatory α-C helix, suggesting the binding of JM could regulate kinase activity. CSPs shown as gradient: Yellow – lower, Red - higher.

**Fig. S2.**
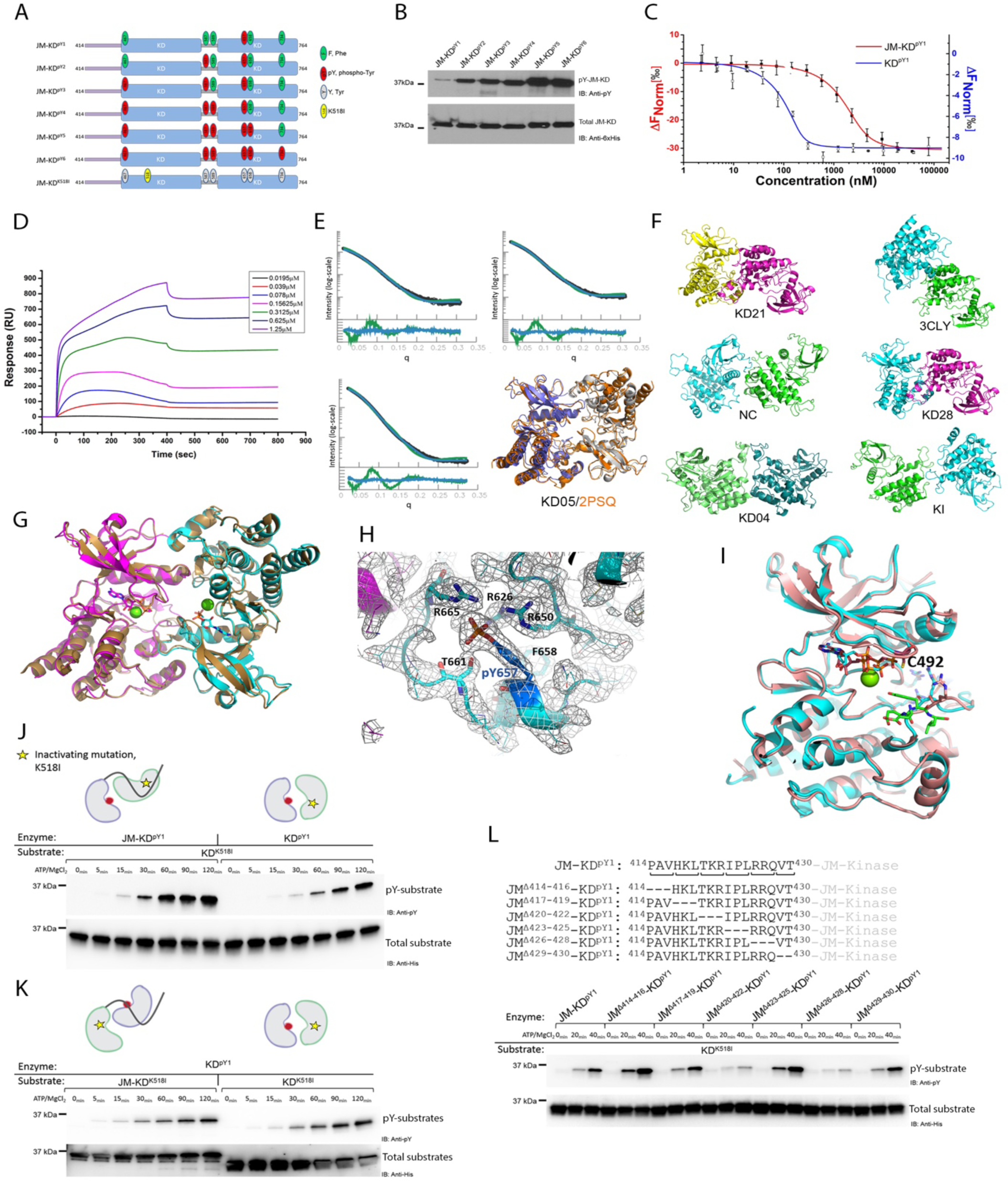
Dimerization of mono-phosphorylated kinase. (**A**) Schematic of JM-KD with progressively phosphorylation KD (FGFR2_414-764_). Six tyrosine residues on KD were mutated to mimic the sequential phosphorylation pattern of KD (JM-KD^pY1^ to JM-KD^pY6^). (**B**) Phosphorylation states of JM-KD^pY1^ to JM-KD^pY6^ were examined using a phosphotyrosine pY99 antibody. An anti-6xHis tag antibody was used to probe for total proteins as the loading control. (**C**) The ‘apparent’ dimerization K_d_ of JM-KD^pY1^ (red) and KD^pY1^ (blue) determined using MST. JM-KD^pY1^ and KD^pY1^were labelled with Atto488 dye then titrated with unlabelled JM-KD^pY1^ and KD^pY1^. (**D**) Dimerization of JM-KD^pY1^ was examined using surface plasmon resonance (SPR). JM-KD^pY1^ was immobilised on a CM4 chip by amine coupling, a serial dilution of JM-KD^pY1^ was injected for 400 seconds and washed with buffer for further 400 seconds. The binding affinity was calculated using steady-state fitting model. (**E**) SAXS description of the structure of KD^pY1^. Top panel: the calculated SAXS pattern for 1-state (green) and 3-state (blue) models fitted to the experimental data (black) at different total maximum KD^pY1^ concentrations (top left plot, 65 µM; top right, 130 µM, and bottom 210 µM). For each scattering dataset the bottom panel shows the residuals of fit for 1-state (green) and 3-state models (blue). The resolution, q, is given in 1/Å and the intensity is given in arbitrary units. The structures in the bottom right panel show that the best-fitting crystal structure-derived (orange), and *in silico*-docked (blue/grey KD05) represent the same conformation. (**F**) Additional SAXS structures relevant to fitting of KD^pY1^ or JM-KD^pY1^ to SAXS data, reported in Fig. 2d, e and Table S3.(**G**) Phosphorylated chains A (magenta) and C (cyan) superimposed onto chains C and D (grey) together with both chains from the unphosphorylated FGFR2 kinase structure 2PSQ. ATP shown as a stick model and Mg^2+^ as green spheres. (**H**) A-loop (chain B) in its 2FoFc electron density. (**I**) Superimposition of KD^pY1^ (chain B) with 2PVF (salmon). (**J**) At the basal state, the presence of JM in the substrate-acting molecule (JM-KD^K518I^) is required for the recruitment of kinase-acting molecule (KD^pY1^). This asymmetric dimer configuration is required for the enhancement of transphosphorylation as the phosphorylation levels of substrates (left panel: JM-KD^K518I^, right panel: KD^K518I^) were examined using a phosphotyrosine antibody (pY99). An anti-6xHis tag antibody was used to probe total proteins as the loading control. (**K**) The JM from JM-KD^pY1^ cannot recruit and phosphorylate KD^K518I^. figs. S2J and S2K demonstrate a role of kinase activation at basal state by which the JM interacts *in trans*, recruiting and phosphorylating a substrate molecule. (**L**) Deletion in the JM identifies the critical motifs, ^420^TKR^422^ and ^426^RRQ^428^, for JM to recruit and phosphorylate substrate.

**Fig. S3.**
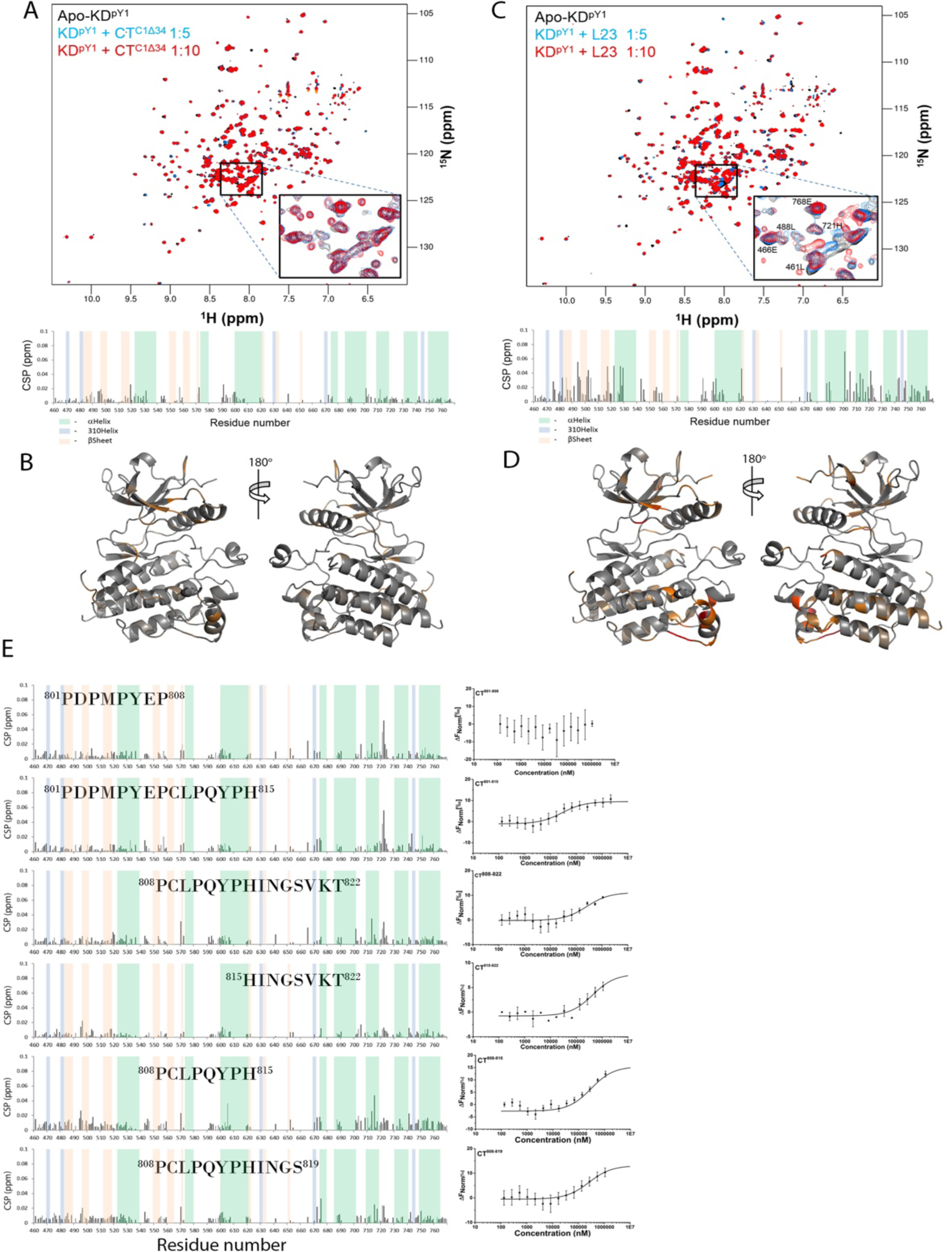
CT^C1^ fragments bind to KD^pY1^. (**A**) The first 24 residues (CT^C1Δ34^) of CT shows negligible interaction with^15^N-labelled KD^pY1^. There is no significant CSPs observed in the HSQC spectra and calculated CSPs. (**B**) Mapping of the weak CSPs from CT^C1Δ34^ binding on the KD^pY1^ crystal structure. (**C**) The last 23 residues (Last 23, L23) of CT shows major interaction with ^15^N-labelled KD^pY1^. Upon L23 binding, CSPs were observed in the HSQC spectra and calculated CSPs shows two clusters of residues can respond to L23 peptide binding. (**D**) Mapping of the CSPs from L23 binding on the KD^pY1^ crystal structure. (**E**) Using small CT peptides (801-808, 801-815, 808-822, 815-822, 808-815, and 808-819. Sequences are shown in the Figure) for HSQC titration with ^15^N-labelled KD^pY1^ in order to narrow down sequence-specific binding region on kinase. The binding of each peptide to KD^pY1^ was also confirmed using MST as shown on the right.

**Fig. S4.**
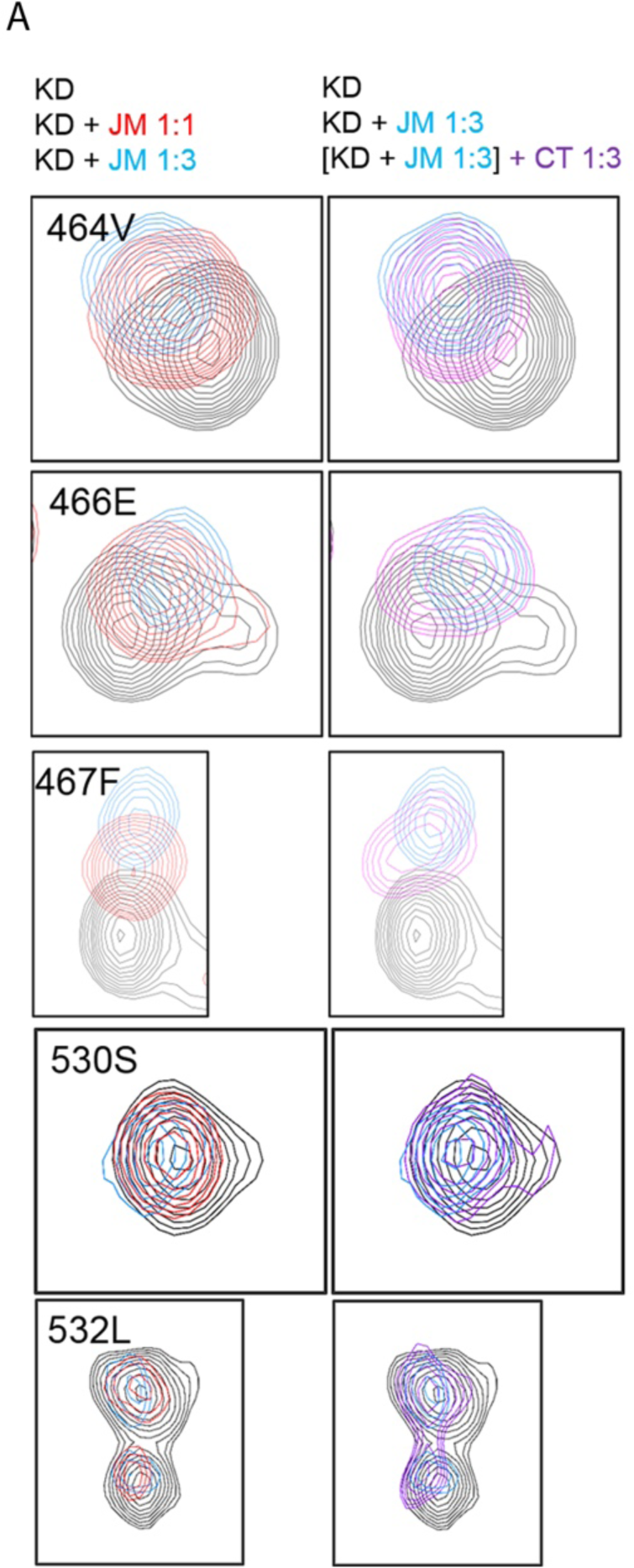
Binding of CT does not compete with JM for binding to KD^pY1^. (**A**) ^15^N-labelled KD^pY1^ was titrated with 1:1 and 1:3 ratio of JM first. CSPs of selected residues were shown (the left hand panel. Apo, black; 1:1, red; 1:3, blue). The addition of 1:3 ratio of CT into the preformed JM-^15^N-labelled KD^pY1^ shows the reduction of CSPs (Right panel; Apo, black; 1 x KD : 3 x JM, blue; [1 x KD : 3 x JM]+3 x CT, purple).

**Fig. S5.**
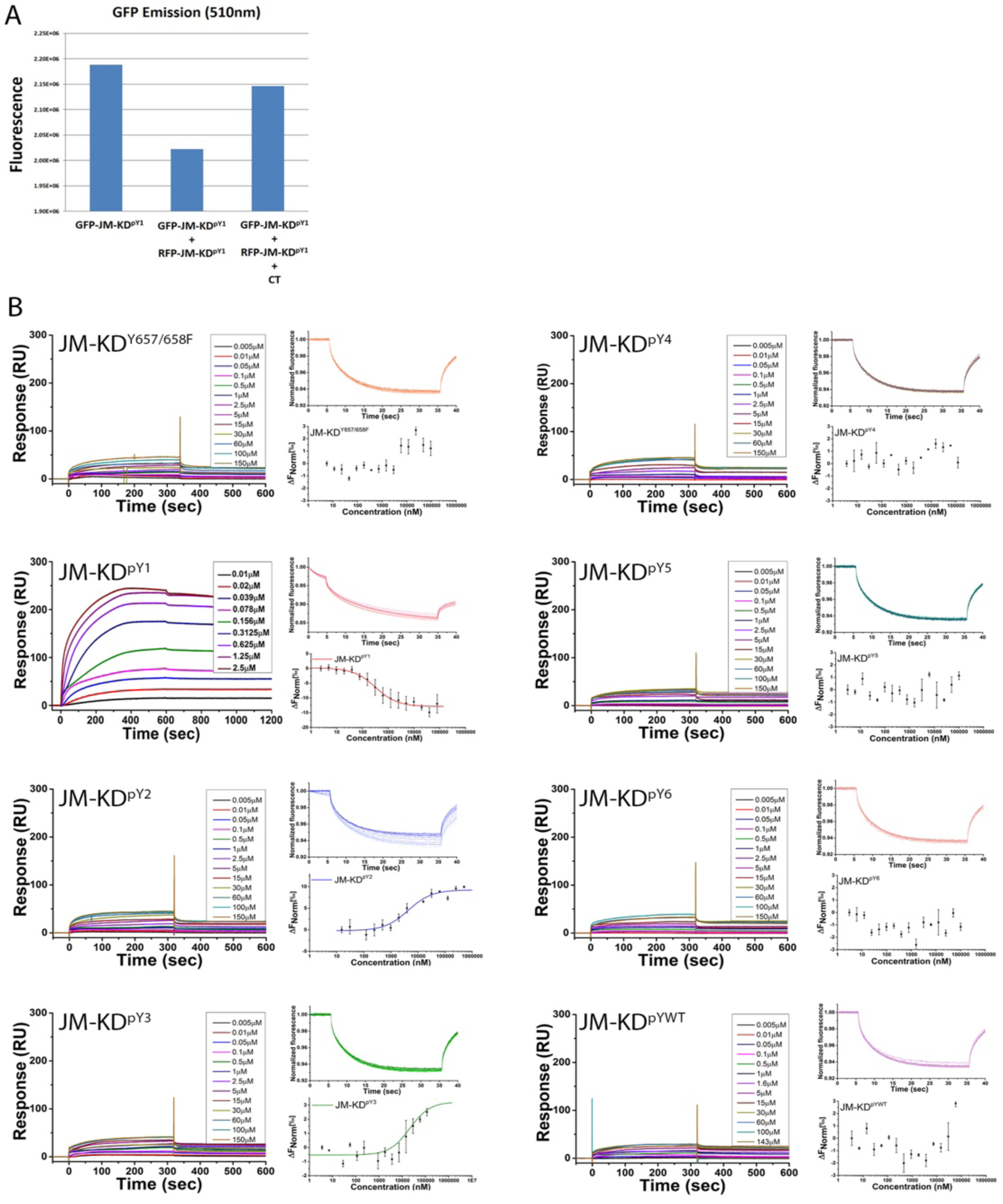
Phosphorylation states control KD and CT^C1^ interaction. (**A**) Steady-state FRET study using GFP and RFP tagged JM-KD^pY1^ demonstrates that the dimer formation of GFP and RFP tagged JM-KD^pY1^ as indicated by the decrease of FRET donor emission (510nm). In addition, the binding of CT to JM-KD^pY1^ results in the dissociation of the asymmetric JM-KD^pY1^ dimer as indicated by the recovery of FRET donor emission (510nm). (**B**) The phosphorylation levels of JM-KD tightly control the interaction with CT. For the SPR experiments, untagged CT (the last 58 residues of FGFR2) was immobilised on a CM4 chip via amine coupling. Kinases with different phosphorylation level (JM-KD^pY1^ to JM-KD^pY6^, and JM-KD^pYWT^, and JM-KD^Y657/658F^) were injected followed by a buffer wash. The binding affinities were determined using stead-state fitting. For the MST measurements, untagged CT was labelled by Atto 488. Two-fold serial dilutions of kinase proteins as described above were used to mix with labelled CT (100nM) and the binding affinities were determined. Both SPR and MST experiments show that the mono-phosphorylated JM-KD is the strongest binding partner for the CT. These experiments provide direct evidence of KD-CT interaction at the basal state.

**Fig. S6.**
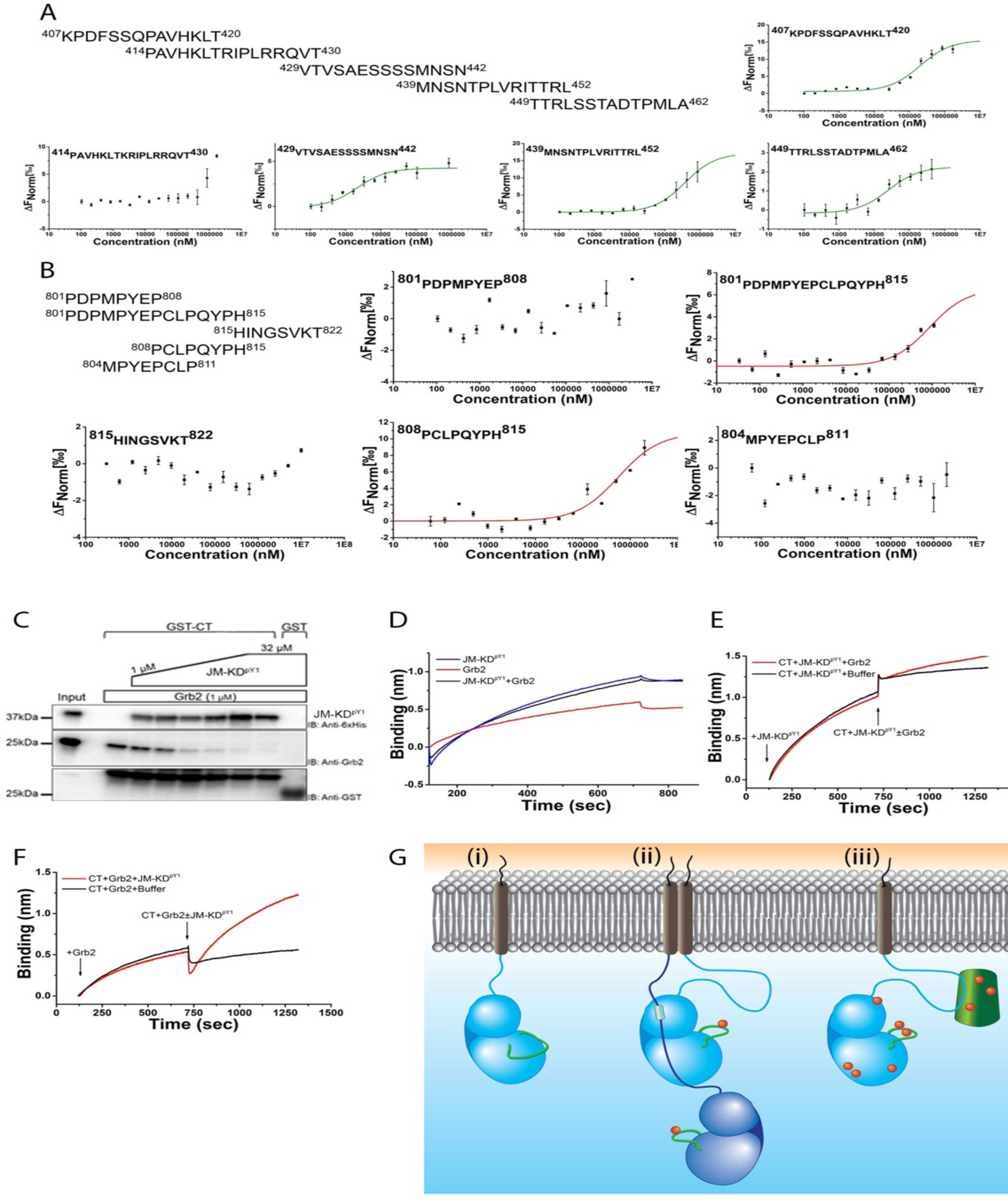
CT-JM interactions and CT competes with GRB2 for binding to JM-KD^pY1^. (**A**) Small JM peptides (residues 407-420, 414-430, 429-442, 439-452, and 449-462) were used for the MST measurements with CT. CT was labelled with Atto 488 and 2-fold serial dilutions of small JM peptides were mixed with 100 nM labelled CT. (**B**) Small CT peptides (residues 801-808, 801-815, 815-822, 808-815, and 804-811) were used for the MST measurements with JM. JM was labelled with Atto488 and 2-fold serial dilutions of small CT peptides were mixed with 100 nM labelled JM. (**C**) GST-CT was used to pulldown GRB2 (1 µM) alone or GRB2 (1 µM) mixed with increased concentrations of JM-KD^pY1^ (1 µM to 32 µM). In the absence of JM-KD^pY1^, GRB2 can be recruited to CT. However, binding of JM-KD^pY1^ to CT competes with GRB2 binding to CT as less Grb2 binds to CT in the presence of JM-KD^pY1^ in a concentration-dependent manner. (**D**) BLI sensors captured with CT show different binding behaviours upon 10µM of JM-KD^pY1^ (black) and GRB2 (red) binding. However, the binding curve for the JM-KD^pY1^-GRB2 mixture shows similar curve (blue) to that of JM-KD^pY1^ alone (black), suggesting that in the presence of JM-KD^pY1^ GRB2 cannot bind to CT as the CT-JM-KD^pY1^ complex is preferable. (**E**) GRB2 does not compete with JM-KD^pY1^ for binding to CT. CT was captured on a BLI sensor and exposed JM-KD^pY1^ at 125 s (arrow) for 625 s, the level of bound JM-KD^pY1^ (10 µM solution in the sample well) on CT was shown as the increase of Binding signal (nm). After adding GRB2 (10 µM solution in the sample well) at 750 s as indicated by an arrow, no significant changes can be observed compared with buffer control (Grb2 addition, red; buffer addition, black). (**F**) JM-KD^pY1^ competes with GRB2 for binding to CT. CT was captured on a BLI sensor and exposed GRB2 at 125 s (arrow) for 625 s, the level of bound GRB2 (10 µM solution in the sample well) on the CT was shown as the increase of Binding signal (nm). After adding JM-KD^pY1^ (10 µM solution in the sample well) at 750 s as indicated by the arrow, a significant drop in the signal following by a graduate increase in signal compared with buffer control (JM-KD^pY1^ addition, red; buffer addition, black) indicating that JM-KD^pY1^ competes with GRB2 for binding to CT. (**G**) Schematic representation of activation of FGFR2^C3^ K*sam* mutant. i: In the absence of stimulation the unphosphorylated FGFR2^C3^ (light blue, JM light blue line) can exist as a monomer freely diffusing through the plasma membrane. ii: Under normal expression levels in non-stimulated cells FGFR2 will self-associate through random collision. Such collision between FGFR2^C3^ molecules in the absence of negative control of CT results in formation A-loop phosphorylation (red spot on green line) of JM latch and asymmetric dimerization to form active enzyme. (Enzyme-like receptor: dark blue, substrate-like receptor: light blue). iii) Active asymmetric dimerization leads to trans-autophosphorylation (red spots) of FGFR2^C3^ which is unrestrained by CT. The conformational change associated with phosphorylation of the KD enables recruitment and phosphorylation of FRS2 (green shape) and recruitment of downstream signalling effector proteins to initiate signal transduction.

**Table S1.** Biophysical measurements of FGFR2 JM, KD, and CT interactions.

| **FGFR2-derived polypeptides binding to JM ($K_d$) measured by MST** |  |
| --- | --- |
| KD<sup>K518I</sup> | 13.4±0.994 μM |
| KD<sup>pY1</sup> | 2.51±0.203 μM |
| KD<sup>pY2</sup> | 6.98±0.877 μM |
| KD<sup>pY4</sup> | 10.2±1.33 μM |
| KD<sup>pY6</sup> | 22.5±1.8 μM |
| KD<sup>pWT</sup> | 20.5±2.04 μM |
| **JM sequences binding to KD<sup>pY1</sup> ($K_d$) measured by MST** |  |
| <sup>407</sup>juxtamembrane region<sup>462</sup> | 2.51±0.203 μM |
| <sup>407</sup>KPDFSSQPAVHKLT<sup>420</sup> | 105±7.37 μM |
| <sup>414</sup>PAVHKLTKRIPRRQVT<sup>430</sup> | 36.8±6.23 μM |
| <sup>429</sup>VTVSAESSSSMNSN<sup>442</sup> | > 1000 μM |
| <sup>439</sup>MNSNTPLVRITTRL<sup>452</sup> | No binding |
| <sup>449</sup>TTRLSSSTADTPMLA<sup>462</sup> | No binding |
| **CT sequences binding to KD<sup>pY1</sup> ($K_d$) measured by MST** |  |
| <sup>801</sup>PDPMPYEP<sup>808</sup> | No binding |
| <sup>801</sup>PDPMPYEPCLPQYPH<sup>815</sup> | 25.8±5.4 μM |
| <sup>808</sup>PCLPQYPHINGSVKT<sup>822</sup> | 257±39.2 μM |
| <sup>815</sup>HINGSVKT<sup>822</sup> | 426±42.5 μM |
| <sup>808</sup>PCLPQYPH<sup>815</sup> | 291±21.4 μM |
| <sup>808</sup>PCLPQYPHINGS<sup>819</sup> | 255±37.9 μM |

| FGFR2-derived polypeptides binding to CT<sup>C1</sup> (μM) measured by SPR |  | SE(Kd) |
| --- | --- | --- |
| JM-KD<sup>Y657/658F</sup> | 5.657E-3 | 1.9E-3 |
| JM-KD<sup>pY1</sup> | 1.652E-4 | 1.6E-5 |
| JM-KD<sup>pY2</sup> | 4.075E-3 | 5.2E-4 |
| JM-KD<sup>pY3</sup> | 3.266E-3 | 8.2E-4 |
| JM-KD<sup>pY4</sup> | 7.873E-3 | 1.7E-3 |
| JM-KD<sup>pY5</sup> | 1.596E-3 | 2.0E-4 |
| JM-KD<sup>pY6</sup> | 4.632E-3 | 9.1E-4 |
| JM-KD<sup>pWT</sup> | 2.309E-3 | 4.8E-4 |
| JM sequences binding to CT<sup>C1</sup> (K<sub>d</sub>) measured by MST |  |  |
| <sup>407</sup>juxtamembrane region<sup>462</sup> | 20.2±2.92 μM |  |
| <sup>407</sup>KPDFSSQPAVHKLT<sup>420</sup> | 200±13.7 μM |  |
| <sup>414</sup>PAVHKLTKRIPRRQVT<sup>430</sup> | No binding |  |
| <sup>429</sup>VTVSAESSSSMNSN<sup>442</sup> | 3.23±0.326 μM |  |
| <sup>439</sup>MNSNTPLVRITTRL<sup>452</sup> | 372±30.3 μM |  |
| <sup>449</sup>TTRLSSSTADTPMLA<sup>462</sup> | 25.4±3.26 μM |  |
| CT<sup>C1</sup> sequences binding to JM (K<sub>d</sub>) measured by MST |  |  |
| <sup>801</sup>PDPMPYEP<sup>808</sup> | No binding |  |
| <sup>801</sup>PDPMPYEPCLPQYPH<sup>815</sup> | 63.6±6.5 μM |  |
| <sup>815</sup>HINGSVKT<sup>822</sup> | No binding |  |
| <sup>808</sup>PCLPQYPH<sup>815</sup> | 56.6±5.5 μM |  |
| <sup>804</sup>MPYEPCL<sup>P811</sup> | No binding |  |

**Table S2.** Statistic parameters of SAXS experiments for KD^pY1^ and JM-KD^pY1^. Top: Analysis of KD^pY1^. KD04: symmetric dimer, formed through C-lobes. KD05: a docked dimer arrangement corresponding to the dimer observed in PDB 2PSQ. KD21: compact asymmetric dimer from ClusPro2.0; the C-lobe of one kinase binds to N- and C-lobes of the receiver kinase. KD28: compact asymmetric dimer from ClusPro2.0; similar to CP21. 3CLY: asymmetric dimer based on PDB 3CLY (kinase C-tail phosphorylation). Mono: monomer, produced by Swiss-Model based on PDB 2PVF. NC3mer: trimer produced by combining a 2PSQ dimer with a NC dimer. KI3mer: trimer produced by combining a 2PSQ dimer with a KI dimer. The normal script, italics and underlining correspond to results based on using different pools as source for multimers. Normal script: using all monomeric and dimeric models, including in silico docked dimers and aggregates. Italics: using only monomeric and dimeric species present in crystal structures of FGFR1 1Y mutants. Underlined: using only monomeric, dimeric and trimeric species present in crystal structures of FGFR1 1Y mutants. For all Pools, the 3-state models fitted the experimental SAXS data significantly better than single or 2-state models (4-state models did not result in significant improvements and were not included), suggesting that at least 3 species co-exist in solution. Bottom: Analysis of JM-KD^pY1^. JM-KD03: two kinases are loosely connected through their juxtamembrane region. JM-KD04: asymmetric dimer, where the juxtamembrane region of the effector kinase latches onto the receiver kinase. JM-KD13: asymmetric dimer, where the juxtamembrane region of the effector kinase latches onto the receiver kinase. 3CLY: asymmetric dimer based on PDB id 3CLY (kinase C-tail phosphorylation). Mono: monomer, produced by Modeller based on PDB 2PVF. Model structures were selected from a pool including crystal structure-derived and *in silico*-docked dimeric and multimeric assemblies (55 different structures in total).

| **KD<sup>pY1</sup> conc \ multi** | **Single-state**<br>**($\chi^2$ / model)** | **Two-state**<br>**($\chi^2$ / models**<br>**[weight])** | **Three-state**<br>**($\chi^2$ / models**<br>**[weight])** |
| --- | --- | --- | --- |
| 2.5 mg/ml | **2.24** / KD21<br>**12.71** / *2PSQ* | **2.13** / 2CLY [0.410];<br>mono [0.590]<br>**4.21** / *2PSQ* [0.307];<br>*mono* [0.693]<br>**3.94** / <u>KI3mer</u><br>[0.105]; <u>mono</u><br>[0.895] | **1.00** / KD05 [0.246];<br>KD04 [0.266]; mono<br>[0.488]<br>**1.35** / *2PSQ* [0.305];<br>*KI* [0.171];<br>*mono* [0.524]<br>**1.29** / *2PSQ* [0.203];<br><u>NC3mer</u> [0.081];<br><u>mono</u> [0.715] |
| 5.0 mg/ml | **4.18** / KD21<br>**36.12** / *2PSQ* | **4.92** / KD28 [0.613];<br>mono [0.387]<br>**14.81** / *2PSQ*<br>[0.438]; *mono*<br>[0.562]<br>**14.47** / <u>KI3mer</u><br>[0.117]; <u>mono</u><br>[0.883] | **1.76** / KD05 [0.311];<br>KD04 [0.294]; mono<br>[0.396]<br>**2.66** / *2PSQ* [0.392];<br>*KI* [0.184];<br>*mono* [0.424]<br>**2.36** / *2PSQ* [0.276];<br><u>NC3mer</u> [0.088];<br><u>mono</u> [0.635] |
| 8.0 mg/ml | **7.18** / CP21<br>**60.01** / *2PSQ* | **9.74** / CP28 [0.718];<br>mono [0.282]<br>**25.63** / *2PSQ*<br>[0.492]; *mono*<br>[0.508]<br>**26.11** / *2PSQ*<br>[0.492]; <u>mono</u><br>[0.508] | **2.89** / CP05 [0.374];<br>CP04 [0.300]; mono<br>[0.325]<br>**4.71** / *2PSQ* [0.463];<br>*KI* [0.184];<br>*mono* [0.352]<br>**4.26** / *2PSQ* [0.339];<br><u>NC3mer</u> [0.089];<br><u>mono</u> [0.572] |
| **JM-KD<sup>pY1</sup> conc \ multi** | **Single-state**<br>**($\chi^2$ / model)** | **Two-state**<br>**($\chi^2$ / models**<br>**[weight])** | **Three-state**<br>**($\chi^2$ / models**<br>**[weight])** |
| 2.5 mg/ml | **15.30** / JM-KD13 | **2.14** / 3CLY [0.154];<br>mono [0.846] | **1.40** / JM-KD04<br>[0.219]; JM-KD03<br>[0.128]; mono<br>[0.652] |
| 5.0 mg/ml | **48.70** / JM-KD13 | **4.98** / 3CLY [0.153];<br>mono [0.847] | **2.48** / JM-KD04<br>[0.218]; JM-KD03<br>[0.125]; mono<br>[0.658] |
| 8.0 mg/ml | **122.05** / JM-KD13 | **10.78** / 3CLY<br>[0.141]; mono<br>[0.859] | **3.04** / JM-KD04<br>[0.206]; JM-KD03<br>[0.111]; mono<br>[0.684] |

**Table S3.**
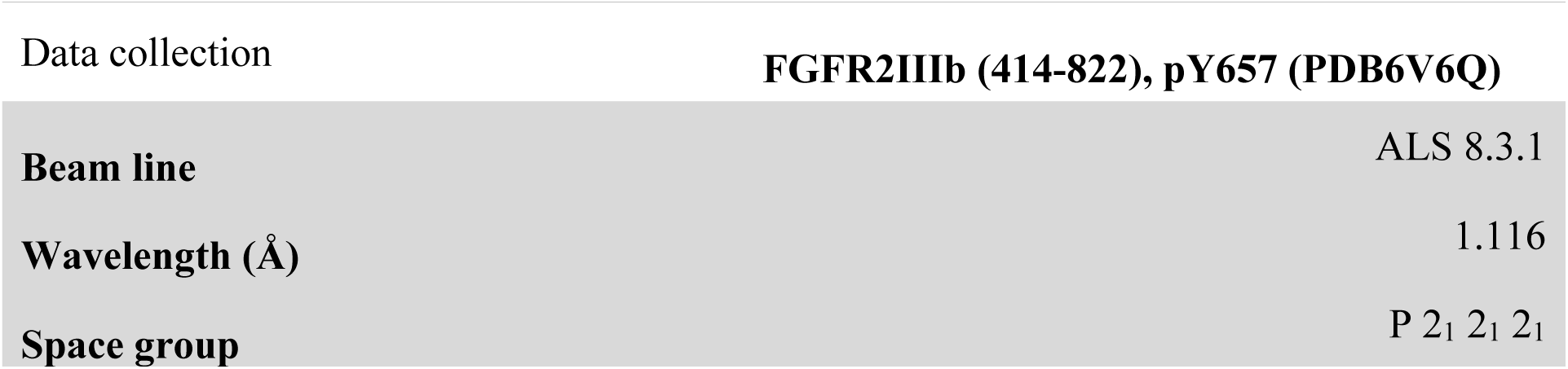
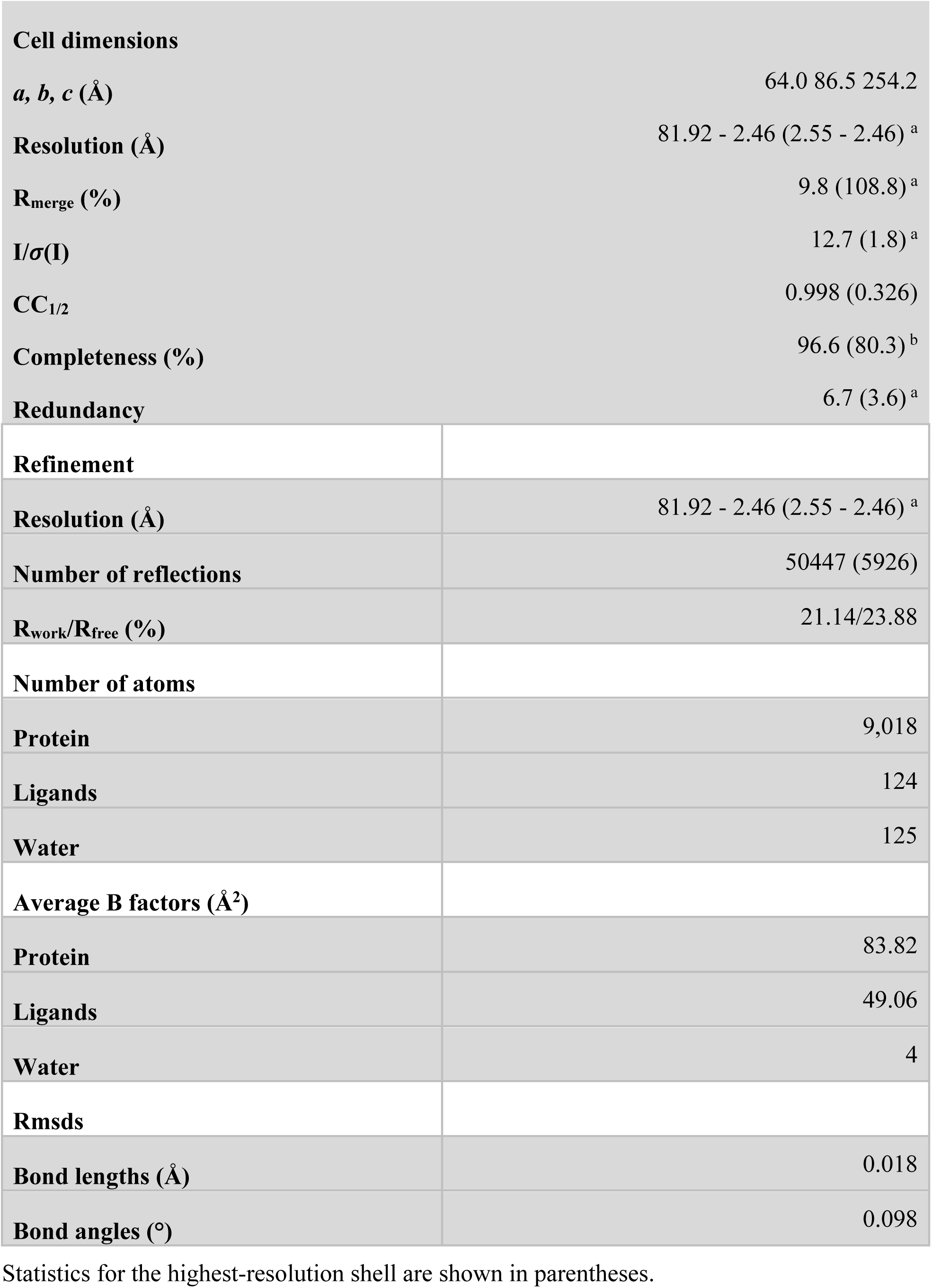
X-ray data collection and refinement statistics. Statistics for the highest-resolution shell are shown in parentheses.

| Data collection |  | FGFR2IIIb (414-822), pY657 (PDB6V6Q) |
| --- | --- | --- |
| Beam line |  | ALS 8.3.1 |
| Wavelength (Å) |  | 1.116 |
| Space group |  | P 2<sub>1</sub> 2<sub>1</sub> 2<sub>1</sub> |

|  |  |
| --- | --- |
| **Cell dimensions** |  |
| *a*, *b*, *c* (Å) | 64.0 86.5 254.2 |
| **Resolution (Å)** | 81.92 - 2.46 (2.55 - 2.46)<sup>a</sup> |
| **R<sub>merge</sub> (%)** | 9.8 (108.8)<sup>a</sup> |
| **I/σ(I)** | 12.7 (1.8)<sup>a</sup> |
| **CC<sub>1/2</sub>** | 0.998 (0.326) |
| **Completeness (%)** | 96.6 (80.3)<sup>b</sup> |
| **Redundancy** | 6.7 (3.6)<sup>a</sup> |
| **Refinement** |  |
| **Resolution (Å)** | 81.92 - 2.46 (2.55 - 2.46)<sup>a</sup> |
| **Number of reflections** | 50447 (5926) |
| **R<sub>work</sub>/R<sub>free</sub> (%)** | 21.14/23.88 |
| **Number of atoms** |  |
| **Protein** | 9,018 |
| **Ligands** | 124 |
| **Water** | 125 |
| **Average B factors (Å<sup>2</sup>)** |  |
| **Protein** | 83.82 |
| **Ligands** | 49.06 |
| **Water** | 4 |
| **Rmsds** |  |
| **Bond lengths (Å)** | 0.018 |
| **Bond angles (°)** | 0.098 |

**Table S4.**
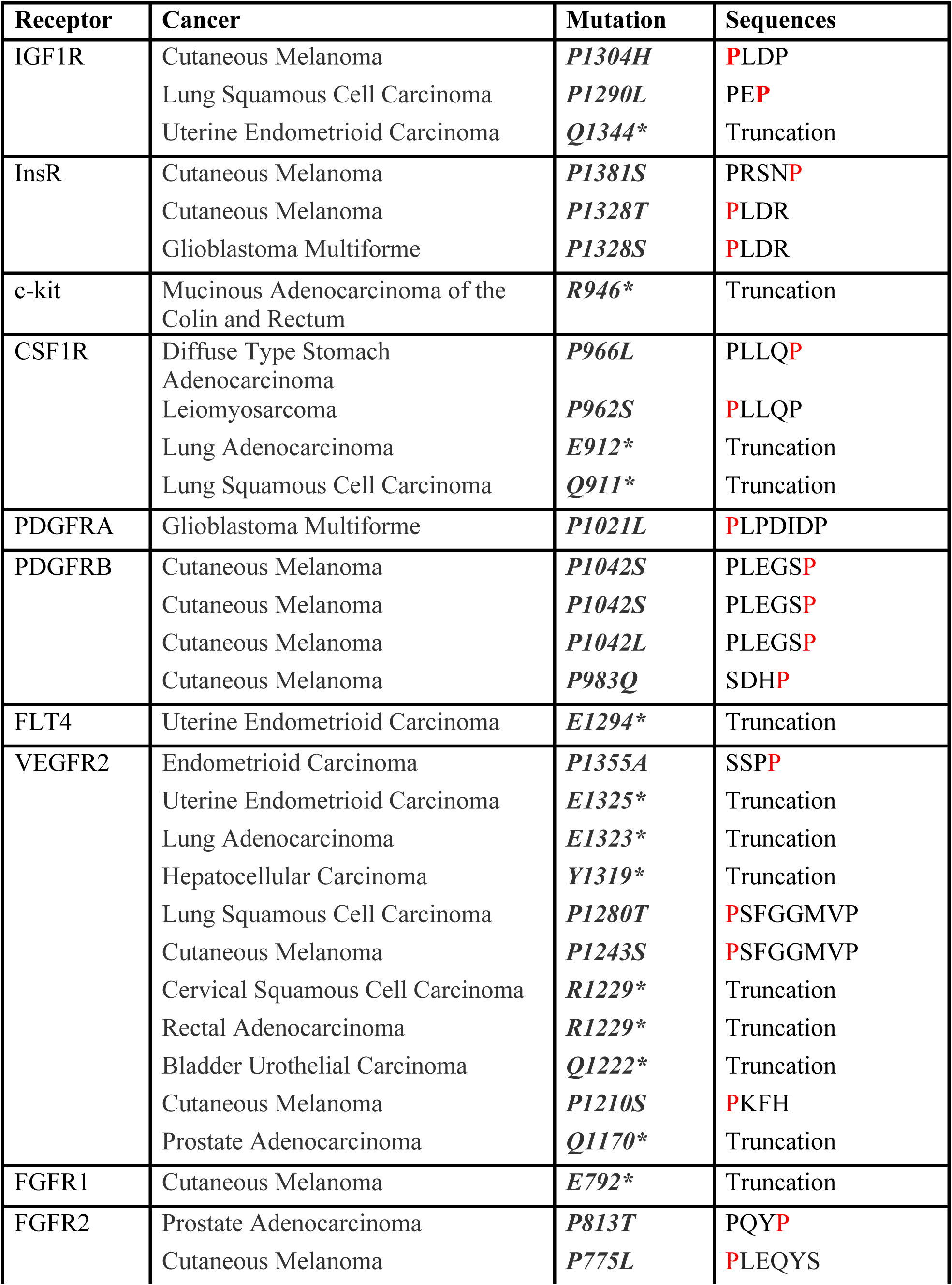
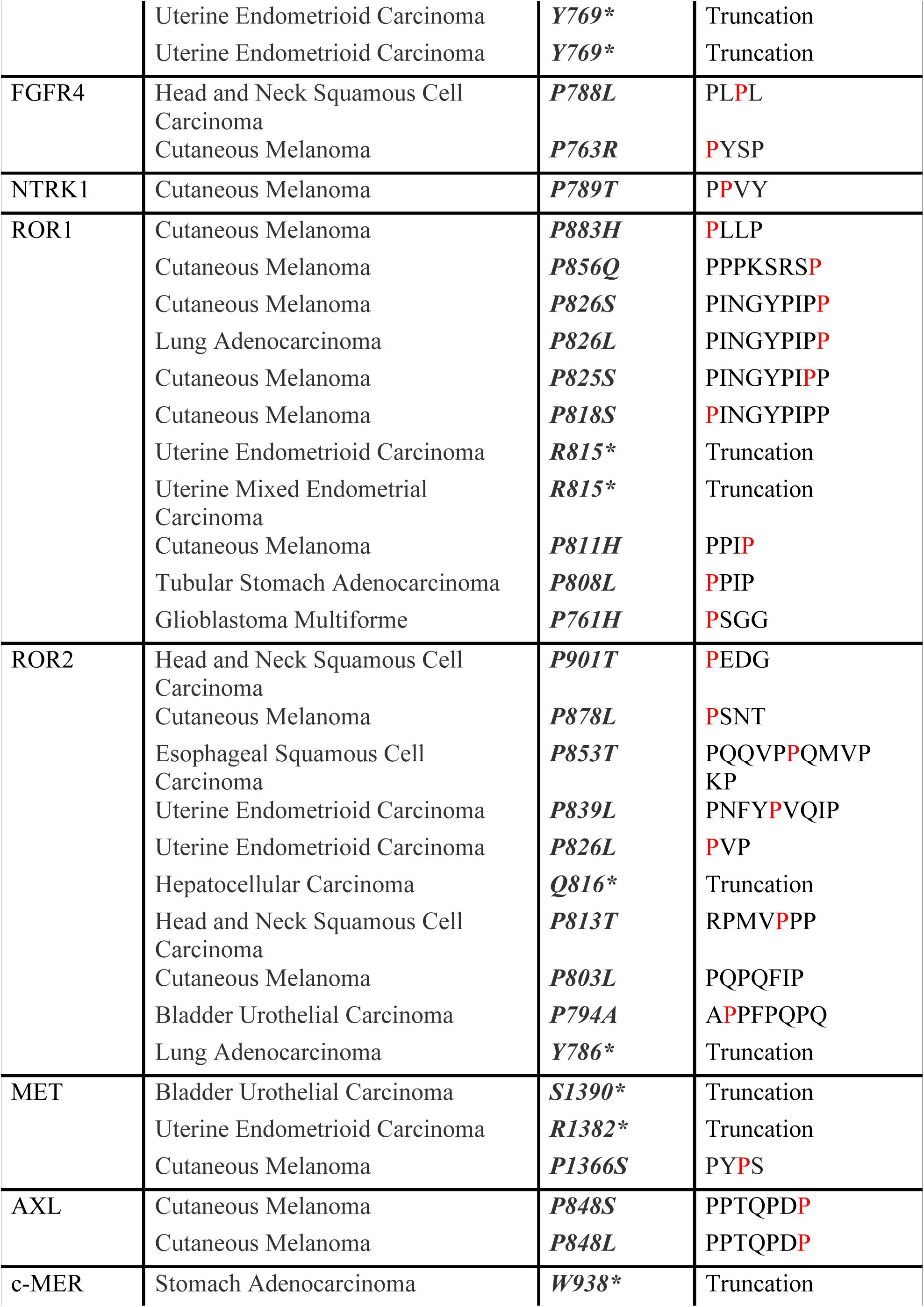
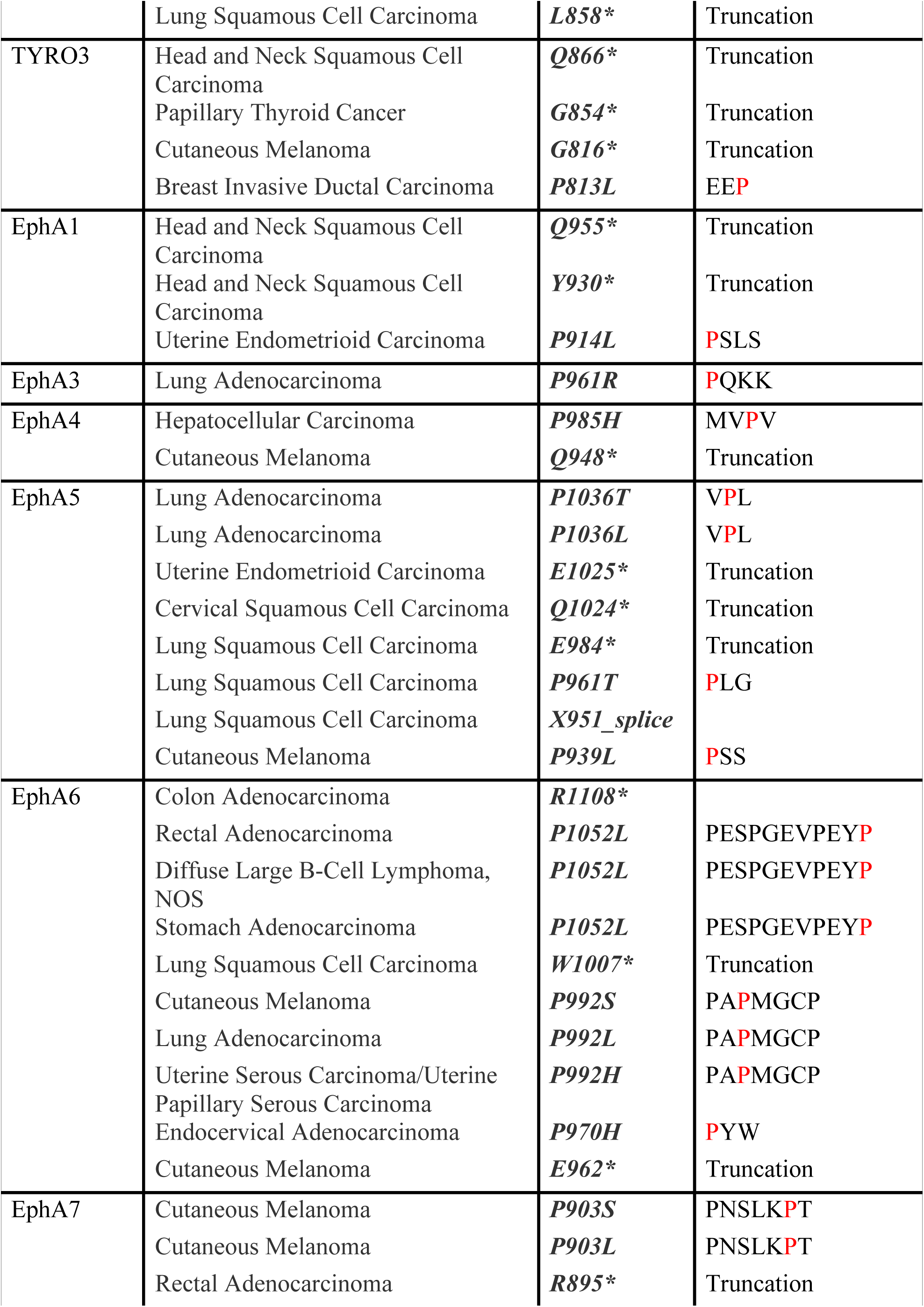
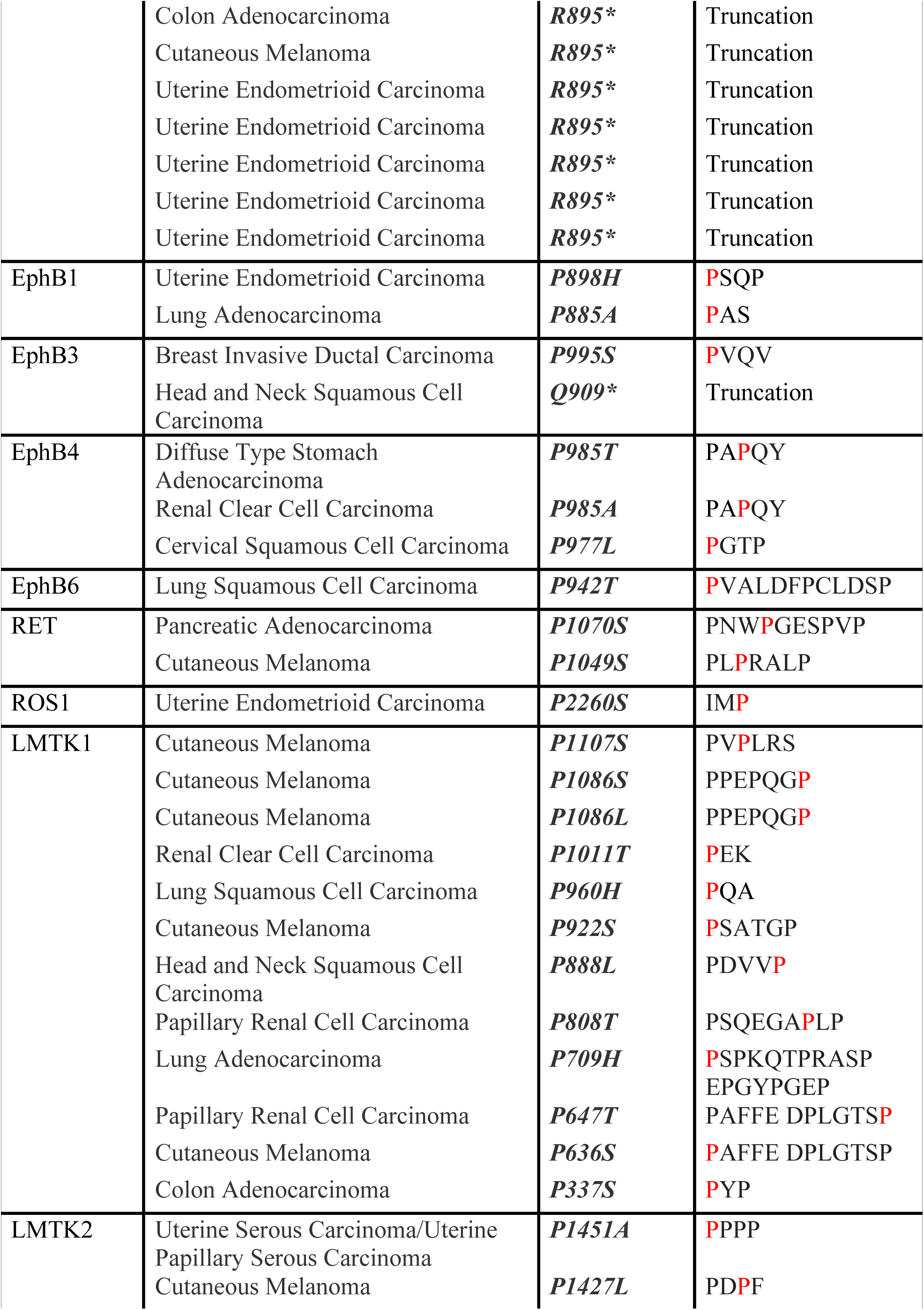
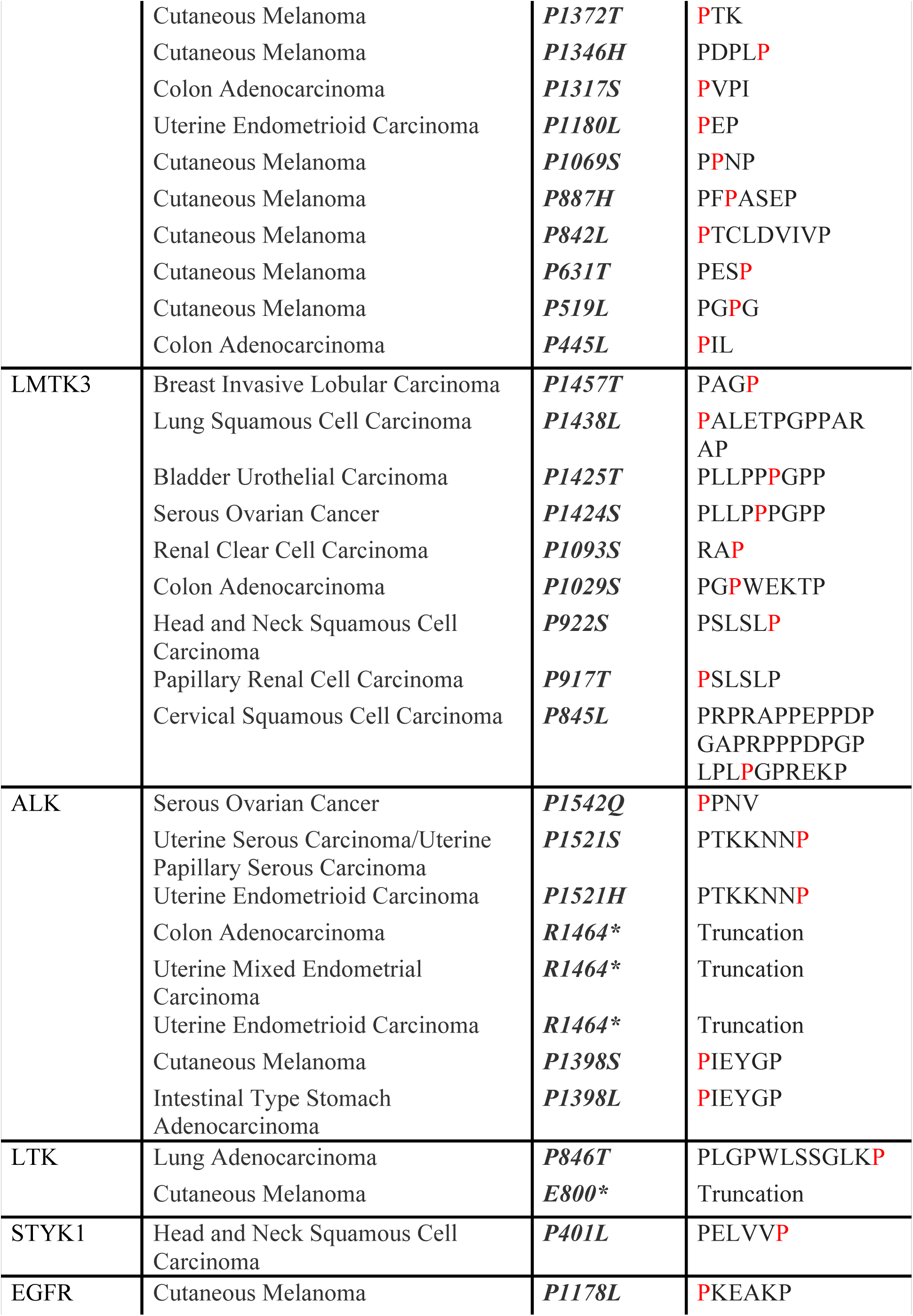
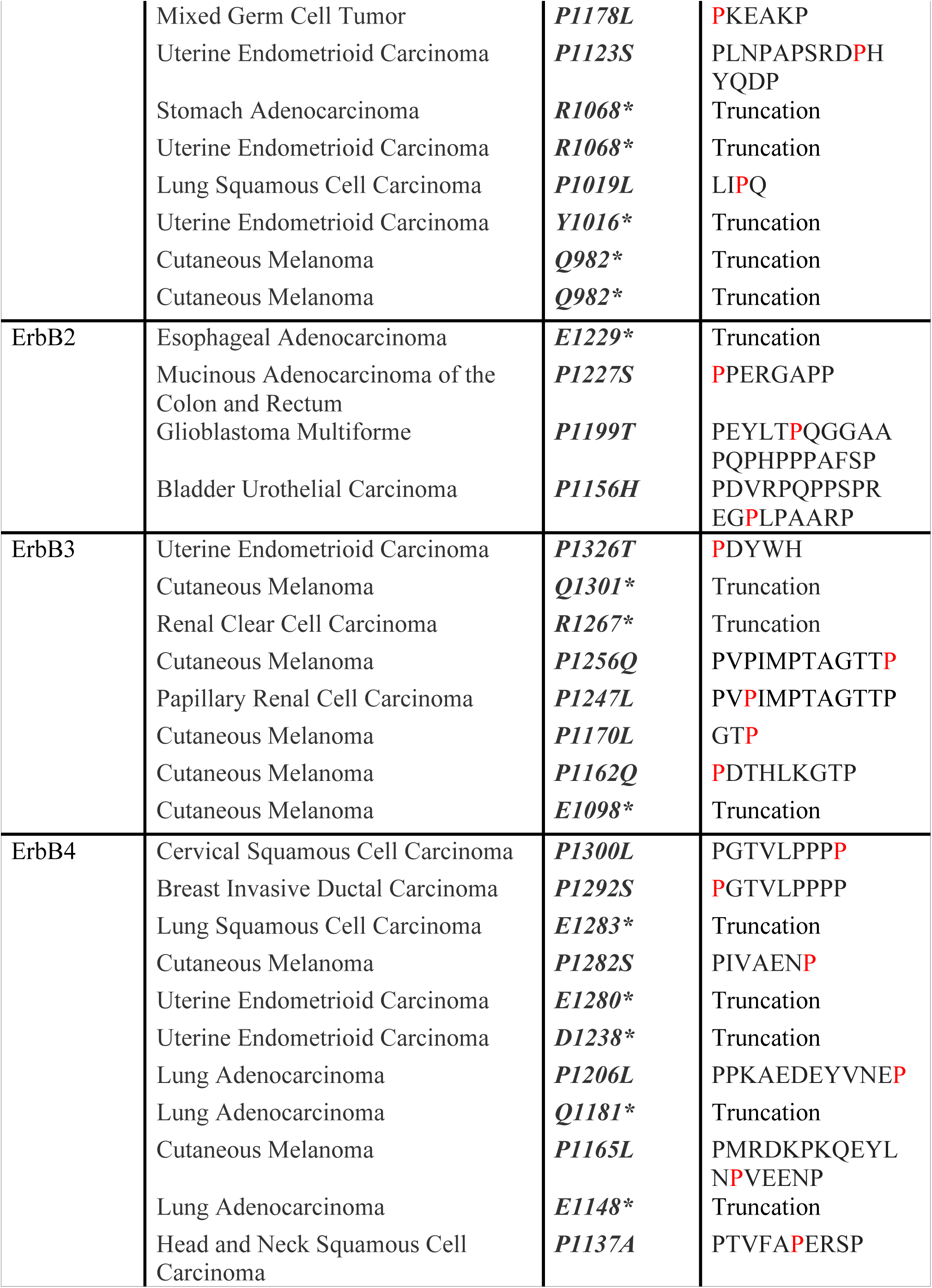
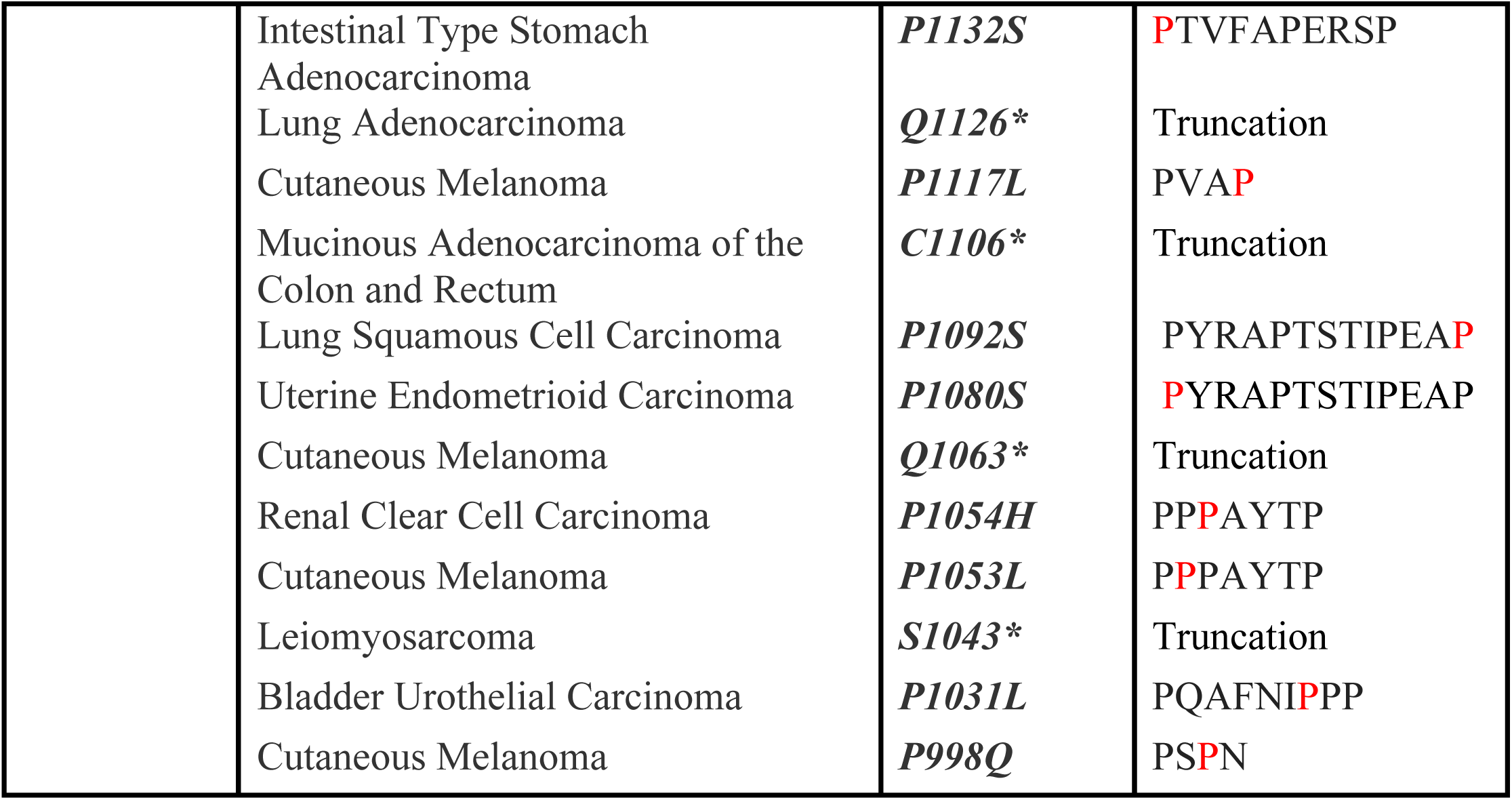
Proline mutants in RTK C-terminal tails and human cancers.

